# Age-Related Alterations in the Expression of Mesencephalic Astrocyte-derived Neurotrophic Factor in the Brain and Their Impact on Neurobehavioral Functions

**DOI:** 10.1101/2025.07.29.667462

**Authors:** Di Hu, Wen Wen, Hui Li, Zuohui Zhang, Hong Lin, Jia Luo

## Abstract

Mesencephalic astrocyte-derived neurotrophic factor (MANF) is a neurotrophic protein localized in the endoplasmic reticulum (ER) and pivotally involved in maintaining ER homeostasis. MANF plays an important role in mitigating neurodegenerative processes. Aging, the primary risk factor for neurodegenerative diseases (NDDs), is associated with significant alterations in ER function. The ER, central to protein synthesis, folding, degradation and secretion (proteostasis), experiences considerable stress in NDDs, which activates the unfolded protein response (UPR). We hypothesized that MANF and UPR is crucial for maintaining proteostasis during aging, but their efficacy declines with age, therefore increasing vulnerability to NDDs. We measured MANF levels in the brain and plasma of 1-, 4-, 11-, and 22-month-old male and female mice. A progressive decline of MANF levels was observed, with the lowest levels detected in 22 months. Reduced MANF expression was found in aged mice across several brain areas, including the cerebral cortex, olfactory bulb, thalamus, hypothalamus, hippocampus, and cerebellum. There was a sex difference in MANF levels in aged mice. Aging also altered the expression of UPR and MANF interacting proteins. Using cerebellar Purkinje cell (PC)-specific MANF deficient mice, we showed that MANF deficiency impaired motor coordination in female, but not male mice. MANF deficiency weakened spatial learning and memory in both male and female mice. Male MANF deficient mice displayed increased sociability, whereas female mice exhibit social withdrawal. Taken together, MANF expression in the brain declined with age and MANF deficiency impacted neurobehaviors in the aging animal in a sex-specific manner.

## 1. Introduction

The endoplasmic reticulum (ER) is a crucial intracellular organelle involved in protein folding and processing, membrane biosynthesis, calcium storage and release, and serves as the entry point of the secretory pathway. Due to these diverse roles, the ER is a key regulator of protein quality control and cellular homeostasis (proteostasis). Disruptions to proteostasis which is triggered by nutrient deprivation, imbalances in calcium or redox levels can lead to the accumulation of misfolded proteins and causes ER stress. ER stress activates the unfolded protein response (UPR), an adaptive mechanism aimed at restoring proteostasis or initiating apoptosis if the damage is irreparable [1]. A significant contributor to aging-related neurodegeneration is the loss of proteostasis and mitochondrial dysfunction [2]. Emerging evidence links ER stress and UPR to the pathogenesis of age-related neurodegenerative diseases such as Alzheimer’s disease (AD), Parkinson’s disease (PD), and Amyotrophic Lateral Sclerosis (ALS) [2, 3]. We hypothesized that the capacity of proteostasis regulation in the brain declines with age, which makes the brain more prone to ER stress and neurodegeneration.

Mesencephalic astrocyte-derived neurotrophic factor (MANF), also known as arginine-rich mutated in early-stage tumors (ARMET), is a 20 kDa secreted protein belonging to a *novel*, evolutionarily conserved neurotrophic factor family [4]. Initially identified as a trophic factor for dopamine neurons, MANF is broadly expressed in the developing and adult central nervous system (CNS) and is upregulated in response to ER stress. MANF deficiency in the brain may activate UPR pathways [5]. Like other components of the UPR, MANF acts to maintain proteostasis under pathological conditions [6, 7]. Beyond this, MANF regulates neurodevelopmental processes including neurogenesis, neuronal survival, differentiation, and neurite outgrowth [8–10]. It also has a neuroprotective role in various neuropathological conditions and reduces neurodegeneration in experimental models of PD, ischemia, spinocerebellar ataxia, and retinal injury by mitigating ER stress [11–16]. Our previous work demonstrated that MANF expression in the brain is developmentally regulated, implying its involvement in neural development [17]. Due to its involvement in the regulation of proteostasis, MANF may also play a role in the process of brain aging.

In the present study, we sought to investigate whether there is an age dependent MANF expression. We measured MANF levels in the brain and plasma of 1-, 4-, 11-, and 22-month-old male and female mice. We demonstrated a progressive decline in both brain and plasma MANF levels with the lowest levels detected in old mice (22 months). There was a sex difference of MANF expression in the plasma and cerebellum; the levels of MANF in old female mice were significantly lower than that of male mice. To mimic MANF scarcity in aged brain, we generated a mouse model of cerebellar Purkinje cells (PC)-specific MANF knockout (KO) to assess the potential involvement of MANF in neurobehavioral functions. MANF KO caused a sex-dependent decline of motor coordination and affected other neurobehavioral parameters. Thus, the progressive reduction of MANF levels in the brain may contribute to aging-associated neurological disorders.

## 2. Materials and Methods

### 2.1 Reagents

Ketamine/xylazine was obtained from Butler Schein Animal Health (Dublin, OH). Fast Tissue/Tail PCR Genotyping Kit (G1001) was obtained from EZ BioResearch (St. Louis, MO). DC protein assay Kit II (5000112) was obtained from Bio-Rad (Hercules, CA). ECL Prime Western Blotting Detection Reagent (45-002-401) was obtained from GE Healthcare Life Sciences (Piscataway, NJ). Anti-ARMET/ARP (MANF) (ab67271), anti-ATF6 (ab203119), anti-phosphorylated IRE1α (ab48187), anti-cleaved caspase-12 (ab62463), anti-TNFα (ab6671) and anti-PDIA1/ P4HB (ab2792) antibodies were from Abcam (Cambridge, MA). Anti-GRP78 (NBP1-06274) antibody was from Novus Biologicals (Littleton, CO). Anti-GRP94 antibody (ADI-SPA-850) was from Enzo Life Sciences (Farmingdale, NY). Anti-β-Actin (3700), anti-PERK (3192), anti-phosphorylated eIF2α (3398), anti-eIF2α (9722), anti-ATF4 (11815), and anti-cleaved caspase-3 (9661) antibodies were from Cell Signaling Technology (Danvers, MA). Anti-XBP1s (658802) antibody was from BioLegend (San Diego, CA). Anti-Iba1 (PA5-27436) and HYOU1 (PA5-27655) antibody was from Thermo Fisher Scientific (Rockford, IL). Anti-MCP1 (AAM-43) antibody was from Bio-Rad (Hercules, CA). Anti-Neuroplastin (AF7818) antibody was from R&D Systems (Minneapolis, MN). Anti-PDIA6 (18233-1-AP) antibody was from Proteintech (Rosemont, IL). Anti-Calbindin (C9848) antibody was from Sigma-Aldrich (St. Louis, MO). HRP-conjugated anti-rabbit (GENA934) and anti-mouse (GENA931) secondary antibodies were from GE Healthcare Life Sciences (Piscataway, NJ). Alexa-488 conjugated anti-mouse (A21202) and Alexa-594 conjugated anti-rabbit (A11012) secondary antibodies were from Life Technologies (Carlsbad, CA).

### 2.2. Animals

Male and female C57BL/6J mice (JAX #000664) of 1, 4, 11, and 22-month-old were obtained from the Jackson Laboratory. Mice were housed under controlled environmental conditions with a 12-hour light/12-hour dark cycle and had *ad libitum* access to food and water in the University of Iowa’s Laboratory of Animal Resources. All experimental animal procedures were approved by the Institutional Animal Care and Use Committee (IACUC) at the University of Iowa (#0032295) and performed following regulations for the Care and Use of Laboratory Animals set forth by the National Institutes of Health (NIH) Guide. The Purkinje cell (PC)-specific MANF knockout (KO) mice was generated and described previously [18]. Briefly, *Manf* ^fl^/^fl^ transgenic mice generated previously [19] were crossed with PC-specific Cre mice B6.Cg-Tg(Pcp2-cre)3555Jdhu/J (purchased from The Jackson Laboratory) to generate *Manf* ^fl/fl^; *Pcp2-Cre*^+/-^ PC-specific MANF KO mice and Cre negative littermate controls that were *Manf* ^fl/fl^; *Pcp2-Cre*^-/-^.

### 2.3. Immunoblotting (IB)

Mice were deeply anesthetized with ketamine/xylazine and perfused transcardially with ice-cold PBS. Brains were promptly removed, bisected sagittally, and the neocortex and cerebellum from the right hemisphere were dissected and stored at −80°C for subsequent analyses. Frozen tissues were incubated in RIPA buffer supplemented with a Protease Inhibitor Cocktail, sodium orthovanadate, and PMSF (Thermo Fisher Scientific, Rockford, IL), and homogenized using an ultrasonic processor on ice. Homogenates were vortexed, followed by centrifugation at 20,000 × *g* for 15 minutes at 4°C. Supernatants were collected, and protein concentrations were quantified using the DC Protein Assay Kit. Equal amounts of protein (30 µg per lane) were separated on 12% SDS-PAGE gels and transferred to PVDF or nitrocellulose membranes. Membranes were blocked for 1 hour at room temperature with 5% BSA in TBS-T (0.1% Tween-20 in TBS). Blots were incubated overnight at 4°C with the following primary antibodies: anti-MANF (1:1000), anti-phospho-IRE1α (1:1000), anti-PDIA1/P4HB (1:1000), anti-cleaved caspase-12 (1:1000), anti-TNFα (1:1000), anti-β-Actin (1:10000), anti-ATF4 (1:1000), anti-PERK (1:1000), anti-cleaved caspase-3 (1:1000), anti-eIF2α (1:1000), and anti-phospho-eIF2α (1:1000), anti-XBP1s (1:1000), anti-MCP1 (1:1000), anti-PDIA6 (1:1000),anti-GRP94 (1:1000,), anti-Neuroplastin (1:4000), anti-GRP78 (1:1000), anti-Iba1 (1:1000) and anti-HYOU1 (1:1000). After washing three times with TBS-T, membranes were incubated with HRP-conjugated secondary antibodies for 1 hour at room temperature. Signals were developed using Amersham ECL Prime Western Blotting Detection Reagent and visualized with a ChemiDoc Imaging System (Bio-Rad). Band intensities were quantified using Image Lab software (Bio-Rad).

### 2.4. Quantification of plasma MANF by enzyme-linked immunosorbent assay (ELISA)

MANF levels in plasma was measured using an ELISA kit (LifeSpan BioSciences, Inc., Olympia, WA). Approximately 100 μL of blood was collected from the tail vein of each mouse using heparin-coated microhematocrit capillary tubes (Thermo Fisher Scientific). Blood samples were centrifuged at 2,000 × *g* for 10 minutes at 4 °C to isolate plasma. Plasma samples were stored at −80°C until analysis. Before the assay, samples were thawed on ice and diluted according to the manufacturer’s instructions. Plasma MANF levels were measured according to the manufacturer’s instruction.

### 2.5. Immunohistochemistry (IHC) and immunofluorescence (IF)

Mice were deeply anesthetized with ketamine/xylazine and transcardially perfused with ice-cold PBS. The left hemispheres were post-fixed in 4% paraformaldehyde (PFA) in PBS for 48 hours at 4°C, followed by cryoprotection in 30% sucrose in PBS for an additional 48 hours. Tissues were embedded in Tissue-Tek optimal cutting temperature (OCT) compound, and sagittal brain sections (10 µm thick) were prepared using a rotary microtome (Leica) and mounted onto Superfrost Plus microscope slides (VWR, Radnor, PA). For IHC staining, sections were washed twice in PBS for 10 minutes each, then blocked for 1 hour at room temperature with a solution containing 1% bovine serum albumin (BSA), 2% normal goat serum, and 0.1% Triton X-100 in PBS. Following the blocking step, sections were incubated with a primary antibody against MANF (1:400) overnight at 4°C. Negative control sections were processed in parallel without the primary antibody. After primary incubation, sections were washed with PBST (PBS containing 0.1% Tween-20) and treated with 3% hydrogen peroxide and 40% methanol in PBS for 20 minutes to quench endogenous peroxidase activity. After thorough rinsing with PBST, sections were incubated with an HPR-conjugated biotinylated goat anti-rabbit IgG secondary antibody (1:200) for 1 hour at room temperature, followed by a 1-hour incubation with an avidin-biotin-peroxidase complex (ABC, Vector Laboratories). Immunoreactivity was visualized using a DAB peroxidase substrate kit (Vector Laboratories), and slides were imaged using an Olympus BX81 microscope. For quantification of MANF staining intensity, representative images were acquired at 10×magnification under identical exposure settings using the accompanying Olympus imaging software. For IF staining, sections were first washed with PBS and PBST, then blocked for 1 hour at room temperature in a solution containing 1% BSA, 2% normal goat serum, and 0.1% Triton X-100 in PBS. After blocking, sections were incubated with primary antibodies against MANF (1:500) and Calbindin (1:500) overnight at 4°C. Following three washes in PBST, sections were incubated with secondary antibody conjugated to Alexa Fluor 594 (red) or 488 (green) (1:200). After another series of three washes in PBST and PBS, nuclear counterstaining was performed using DAPI. Fluorescent images were captured using an Olympus BX81 microscope.

### 2.6. Real-time RT-PCR

Total RNA was extracted from mouse cortex using the RNeasy Mini Kit (Qiagen, Germantown, MD) according to the manufacturer’s instructions. Briefly, 10–20 mg of tissue was homogenized in Buffer RLT supplemented with 1 M dithiothreitol (DTT) using a TissueLyser for 2 minutes. After centrifugation, the supernatant was mixed with 70% ethanol and loaded onto RNeasy spin columns. Columns were washed sequentially with RW1 and RPE buffers, and RNA was eluted in RNase-free water. Reverse transcription was carried out using the High-Capacity cDNA Reverse Transcription Kit with RNase Inhibitor (Applied Biosystems, Pleasanton, CA). The resulting cDNA was stored at −20°C. Quantitative PCR was conducted using TaqMan™ Universal PCR Master Mix. Gene-specific primers for *Manf* and *18s rRNA* (Applied Biosystems) were mixed with cDNA (1:5) in RNase-free water. Reactions were run in triplicate on a 96-well plate, centrifuged, and analyzed using a QuantStudio 3 Real-Time PCR System (Applied Biosystems). Relative gene expression levels were calculated using the ΔΔCt method, with *18s rRNA* serving as the internal control.

### 2.7. Behavioral tests

#### 2.7.1. Behavioral schedule

We conducted a series of behavioral tests for Purkinje cell (PC)-specific MANF knockout (KO) and control (Ctrl) mice at 10 months of age. Mice were weight measured and habituated before being evaluated over a 22-day test period as below: open field (days 1-2), elevated plus maze (day 3), three-chamber sociability test (day 4), Barnes maze (days 8-13), balance beam (days 15-17), and rotarod test (days 18-22). To minimize olfactory cues, the testing apparatus was wiped with a damp cloth and thoroughly dried between mice whenever possible. All behavioral tests were conducted at the Neural Circuits and Behavior Core (NCBC) at the University of Iowa.

#### 2.7.2. Open Field

To evaluate locomotor and anxiety-like behaviors, mice were placed in the center of a black-walled, white-floored open field arena (16″ × 16.5″ × 12″) under ambient lighting conditions (110–130 lux). Testing sessions lasted 10 minutes and were conducted over two consecutive days. Animal movement was recorded and analyzed using EthoVision XT 17 software (Noldus Information Technology, Wageningen, Netherlands). Total distance traveled and time spent in the center area (25 × 25 cm) were measured.

#### 2.7.3. Elevated plus maze

Anxiety-like behavior was assessed using the elevated plus maze (San Diego Instruments, San Diego, CA). The apparatus consisted of four arms (two open and two closed) arranged in a plus configuration and was elevated 50 cm above the floor, with the open arms illuminated at 215 lux by an 18″ ring light (Inkeltech) mounted around the overhead camera. Each mouse was placed in the central zone, facing one of the closed arms, at the start of a 5-minute session, and activity was recorded. The number of entries and time spent in the open and closed arms were analyzed using EthoVision XT 17 software. Ratios of open arm entries to total entries and time spent in the open arms to the total duration were also calculated.

#### 2.7.4. Three-chamber sociability test

Sociability was evaluated using a custom-built three-chamber apparatus made of matte black plastic. The arena consisted of a central chamber and two side chambers of equal dimensions, connected by small openings that permitted unrestricted movement between compartments. A perforated cylindrical container was placed in each side chamber to confine either a novel mouse or a novel object, allowing limited direct interaction through the holes. The test consisted of two consecutive 10-minute phases. In the habituation phase, both cylinders were empty, and the test mouse was placed in the central chamber and allowed to explore all three chambers freely. In the sociability phase, a novel, age- and sex-matched wild-type C57BL/6 mouse (10 months old) was placed in one cylinder, and a novel inanimate object in the other. Mouse behavior was recorded and analyzed using EthoVision XT 17 software. The time spent in each chamber and the duration of direct interaction with each cylinder were quantified to assess social preference.

#### 2.7.5. Barnes maze

Spatial learning and memory were evaluated using the Barnes maze (San Diego Instruments, San Diego, CA), which consisted of a circular platform (92 cm in diameter) elevated 90 cm above the floor, with 20 equally spaced holes (5 cm in diameter) around the perimeter. One hole served as the escape hole and was connected to an escape box, while the remaining 19 holes were closed. Visual cues were placed on the surrounding walls to facilitate spatial navigation. The maze was brightly illuminated at approximately 1300 lux to promote escape-seeking behavior. On day 0, mice underwent a single 60-second habituation trial. If a mouse failed to locate the escape hole within this time, it was gently guided to the escape box. The escape hole location was not used during subsequent training/acquisition and probe testing sessions. From days 1 to 4, mice underwent four acquisition trials per day (maximum 180 seconds per trial, with a 15-minute intertrial interval). During each trial, primary latency (time to first locate the escape hole) and primary distance traveled (total path length to first reach the escape hole) were recorded using EthoVision XT 17 software. On day 5, a 90-second probe trial was conducted with the escape box removed to assess memory retention. The maze was conceptually divided into four quadrants, each containing five holes. The target quadrant was defined as the one centered on the former location of the escape hole. The percentage of time spent in the target quadrant relative to the total trial duration was calculated to evaluate spatial memory. Searching strategies during acquisition and probe trials were manually recorded and categorized as direct, serial, or random. A direct strategy was defined as the mouse navigating directly to the target hole or to one of the holes immediately adjacent to it. A serial strategy involved the mouse systematically inspecting adjacent holes by traveling through consecutive holes around the periphery of the maze until the escape hole was located. A random strategy was characterized by disorganized movement across the platform, without a consistent pattern, until the escape hole was found.

#### 2.7.6. Balance beam

Motor coordination and balance as evaluated using the Balance Beam Test (MazeEngineers, Conduct Science, Skokie, IL). The apparatus consisted of an 80 cm-long beam elevated 50 cm above a padded surface, with a safe zone containing clean bedding positioned at one end to serve as a goal. Three beam diameters were used: 24 mm, 12 mm, and 6 mm. Mouse behavior during the trials was recorded using both a front-facing and an overhead camera. Mice were trained for two consecutive days, completing three trials per beam width each day in the order of 24 mm, 12 mm, and 6 mm, with 5-10 minutes of rest between trials. On the test day, each mouse again completed three trials per beam width in the same sequence, and the latency to cross each beam was recorded for valid trials. Trials were considered invalid and excluded from latency analysis if the mouse stalled (remained motionless for more than 5 seconds), reversed direction, fell from the beam, or required prompting.

#### 2.7.7. Rotarod test

Motor coordination was assessed using an accelerating rotarod apparatus (IITC Life Science, Woodland Hills, CA). Mice were tested over five consecutive days. Each mouse was gently placed on the rotating rod (initial speed: 4 rpm), facing away from the experimenter and oriented opposite to the direction of rotation to encourage forward locomotion and reduce immediate falls. The rod accelerated from 4 to 40 revolutions over the first 30 seconds and then maintained 40 rpm for the remainder of the 5-minute trial. Each mouse completed three trials per day, with 10–15 minutes of rest between trials. Latency to fall was automatically recorded. Trials in which the mouse exhibited passive rotation-defined as clinging to the rod and rotating without active movement for two full consecutive revolutions-were terminated, and the latency was still recorded. The average latency across the three trials was calculated for each mouse.

### 2.8. Statistical Analysis

All statistical analyses were performed using GraphPad Prism 10.0 (GraphPad Software, La Jolla, CA). Values were mean ± standard error of the mean (SEM). Statistical analyses were conducted using Welch’s *t*-test, one-way or two-way analysis of variance (ANOVA), followed by Tukey’s post hoc test, when appropriate. Pearson correlation analysis was used to assess the relationship between body weight and beam-crossing time on the 12 mm and 6 mm beams in female mice. Fisher’s exact test was used for the Barnes Maze probe test searching strategy. A p value < 0.05 was considered statistically significant.

## 3. Results

### 3.1. Age- and sex-related changes in MANF expression in the brain and plasma

We first examined the expression levels of MANF protein in the cerebral cortex and cerebellum of 1-, 4-, 11-, and 22-month-old mice using immunoblotting (IB) analysis. While the expression of MANF in the male cortex was observed to be the highest at 1 month, this expression gradually decreased with age to significantly low levels by 22 months (Figs. 1A and B). In female cortex, MANF levels at 1 month were also significantly higher as compared to those of 4, 11, and 22 months (Figs. 1C and D). A similar age dependent decline in MANF in the cerebellum was also present in both males and females (Figs 1A–D). We also evaluated the expression levels of MANF mRNA in the cerebral cortex. The maximal MANF mRNA expression was at 4 months, then declined thereafter (Figs 1E and F).

**Figure 1.**
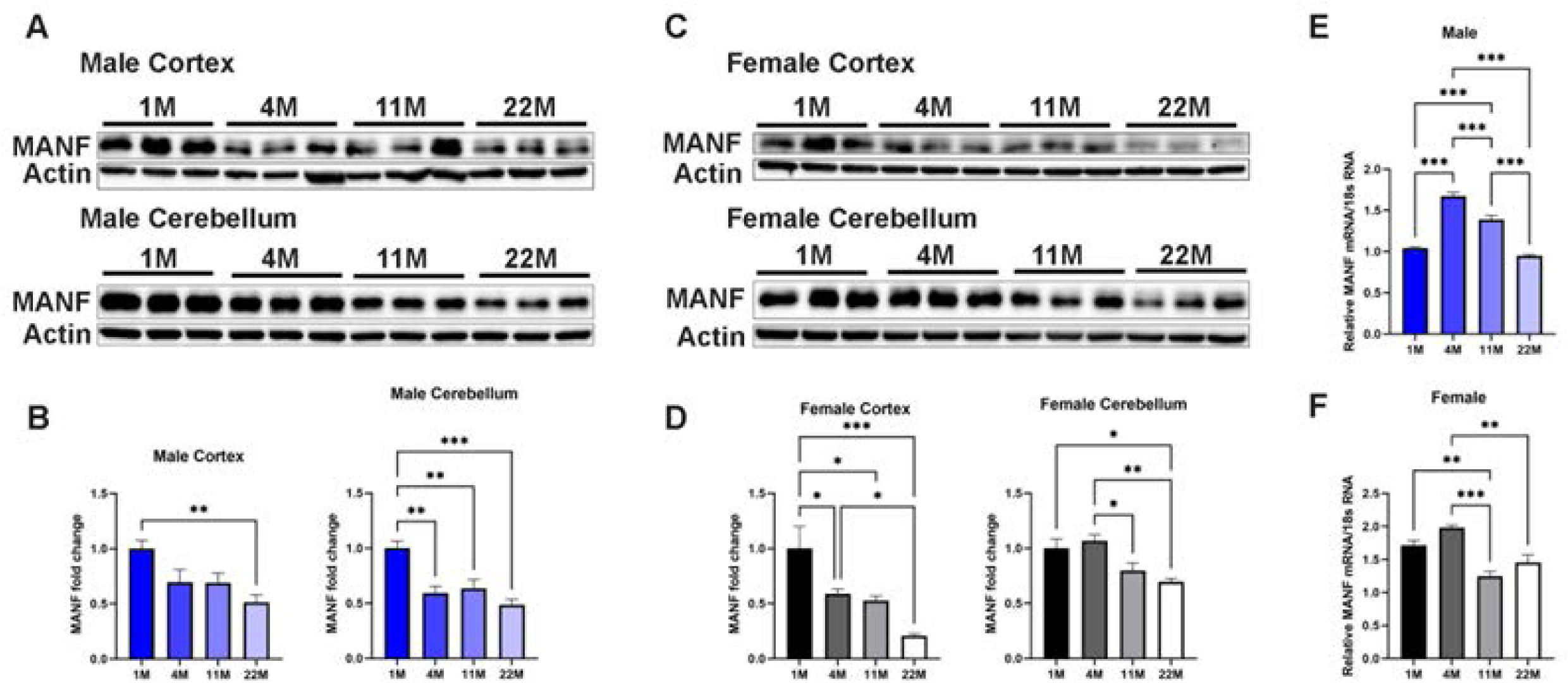
Age-dependent alterations of MANF expression levels in mouse brain. **A** and **C**: Proteins were extracted from the cerebral cortex and cerebellum of 1-, 4-, 11-, and 22- month-old male and female mice. The expression of MANF was examined by immunoblotting (IB) analysis. **B** and **D:** The expression of MANF was quantified as described in the *Materials and Methods* and normalized to the levels of actin. Data were expressed as mean ± SEM, *n* = 5-6 per group. Data were analyzed by One-way ANOVA followed by Tukey’s post hoc test. \**p* < 0.05, \*\**p* < 0.01, \*\*\**p* < 0.001. **E** and **F:** mRNA levels of MANF in the cerebral cortex of 1-, 4-, 11-, and 22- month-old male (E) and female (F) mice were determined by RT-qPCR as described in the *Materials and Methods*. Data were expressed as mean ± SEM, *n* = 6 per group. Data were analyzed by One-way ANOVA followed by Tukey’s post hoc test, \*\**p* < 0.01, \*\*\**p* < 0.001.

We next investigated the levels of MANF protein in the plasma using ELISA. As shown in Fig. 2E, the circulating levels of MANF declined with age and were significantly lower in 11- and 22- month-old male mice relative to 1-month-old male mice. In female mice, compared to 1-month-old mice, MANF plasma drastically decreased after 4 months.

**Figure 2.**
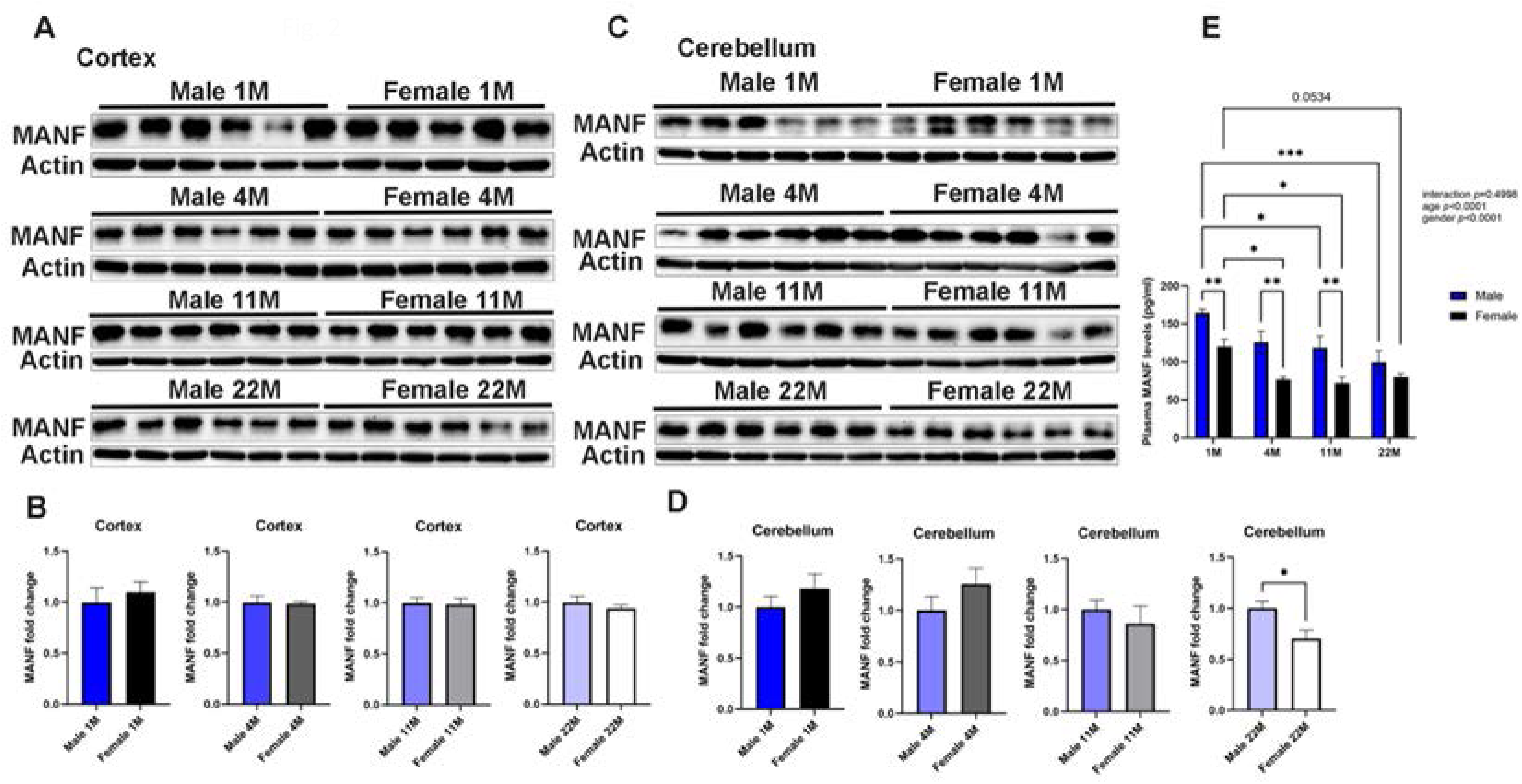
Sex differences of MANF levels in the brain and plasma of mice. **A** and **C**: Proteins were extracted from the cerebral cortex and cerebellum of 1-, 4-, 11-, and 22- month-old male and female mice. The expression of MANF was examined by IB analysis. **B** and **D:** The expression of MANF was quantified as described in the *Materials and Methods* and normalized to the levels of actin. Data were expressed as mean ± SEM, *n* = 5-6 per group. Data were analyzed by Welch’s *t*-test, \**p* < 0.05. **E**. The protein levels of MANF in the plasma of male and female mice at 1-, 4-, 11-, and 22-month-old mice were determined by ELISA. Data were expressed as mean ± SEM, *n* = 6 per group. Data were analyzed by Two-way ANOVA followed by Tukey’s post hoc test, \**p* < 0.05, \*\**p* < 0.01, \*\*\**p* < 0.001.

We sought to determine whether there was a sex difference in MANF expression in the brain and plasma. In the cerebral cortex, there was no difference in MANF expression between male and female mice (Figs. 2A and B). In the cerebellum of 22-month-old mice, the levels of MANF expression in females were significantly lower than males (Figs. 2C and D). In the plasma, levels of MANF in 1-, 4-, and 11-month-old female mice were significantly lower than that of male mice (Fig. 2E). Although its levels in the plasma were lower in 22-month-old female mice than age-matched male mice, the difference did not reach statistical significance (Fig. 2E). These findings indicate that generally MANF levels in the cerebellum and plasma of female mice were less than that of male mice.

We examined the distribution of MANF in the brain of adolescent (one month) and old mice (22 months) using immunohistochemistry (IHC) (Figs. 3 and 4). For male mice, high levels of MANF expression were observed in adolescent brain, particularly the hippocampus, olfactory bulb, cerebral cortex, cerebellum, thalamus and hypothalamus and its expression in these regions was significantly lower in old mice (Fig. 3). It appeared that the levels of MANF in the midbrain were similar between adolescent and old male mice. For female mice, similarly the levels MANF high in these brain regions and but decreased in the olfactory bulb, cerebellum, thalamus and hypothalamus of old mice (Fig. 4). The expression of MANF in the hippocampus, cerebral cortex and midbrain appeared similar between adolescent and old female mice. Although its overall expression in the hippocampus was similar between adolescent and old female mice, MANF levels some specific areas of the hippocampus such as Cornu ammonis (CA)2 and dentate gyrus (DG) declined in old mice (Fig. 4). In the cerebellum, MANF expression in Purkinje cells and granule cells markedly declined with age in both male (Figs. 3B and 3D) and female mice (Figs. 4B and 4D). Taken together, these results indicate that MANF expression declines with age and exhibits sex-specific variation.

**Figure 3.**
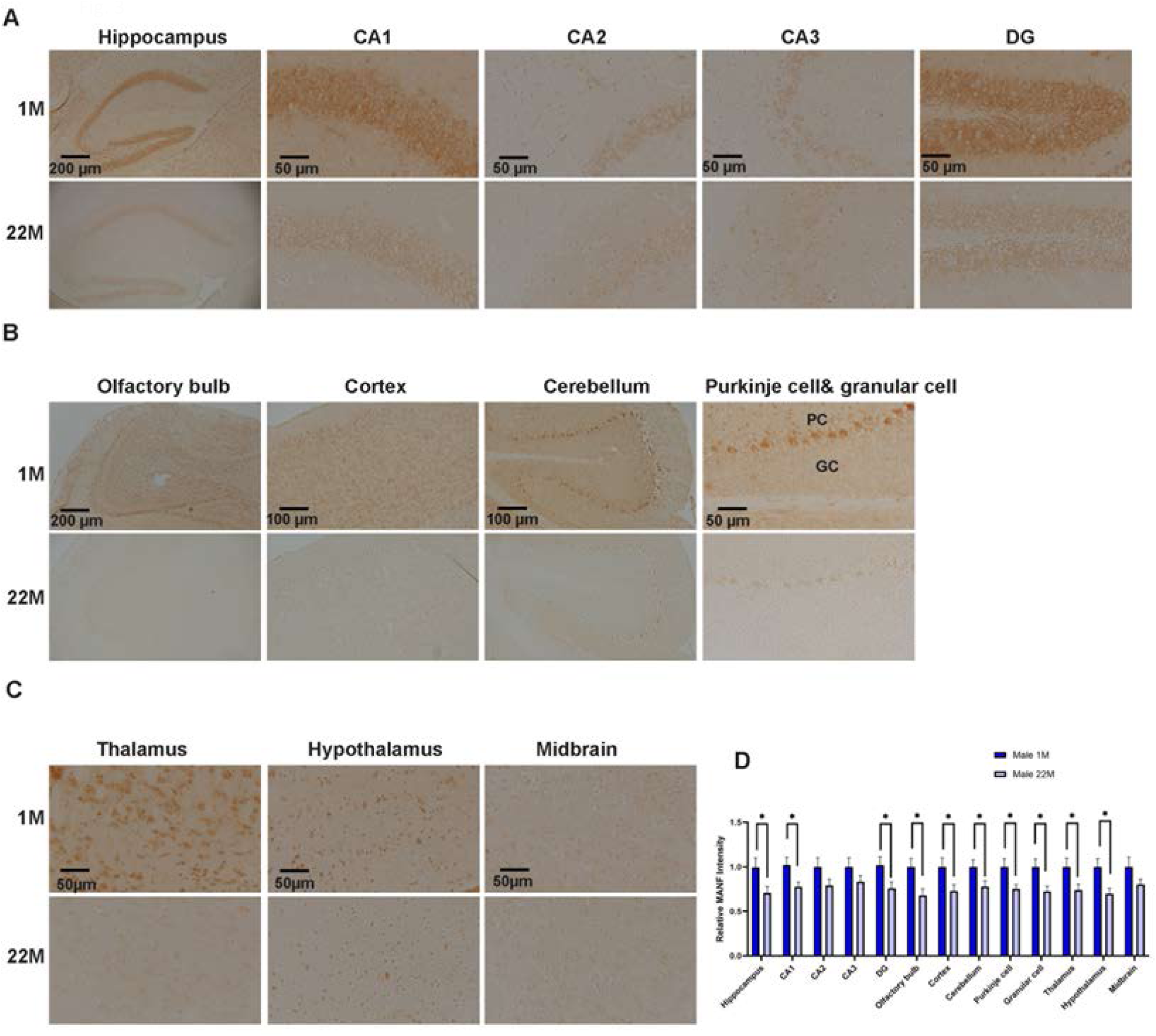
Age-dependent expression of MANF in various brain regions of male mice. **A-C**: The brain tissues obtained from 1- and 22-month-old male mice were fixed and sectioned. The expression of MANF in the hippocampus (CA1, CA2, CA3, and DG), olfactory bulb, cerebral cortex, cerebellum (Purkinje cells (PC) and granular cells (GC)), thalamus, hypothalamus, and midbrain were examined by immunohistochemistry (IHC). **D:** The intensity of MANF in these brain regions was quantified as described in the *Materials and Methods*. Data were expressed as mean ± SEM, *n* = 6 animals per group. Data were analyzed by Welch’s *t*-test, \**p* < 0.05.

**Figure 4:**
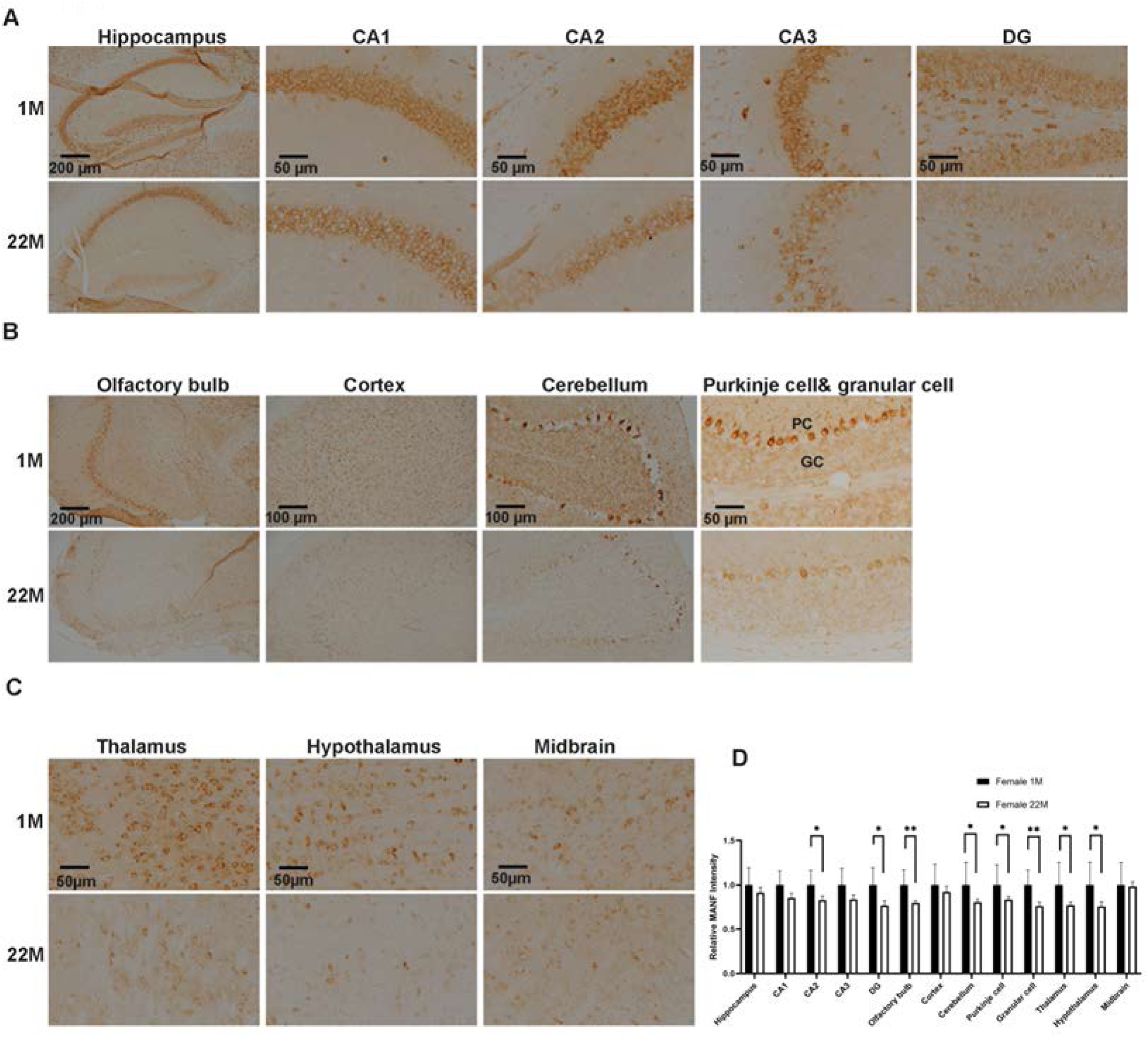
Age-dependent expression of MANF in various brain regions of female mice. **A-C**: The brain tissues obtained from 1- and 22-month-old female mice were fixed and sectioned. The expression of MANF in the hippocampus (CA1, CA2, CA3, and DG), olfactory bulb, cerebral cortex, cerebellum (PC and GC), thalamus, hypothalamus, and midbrain were examined by immunohistochemistry (IHC). **D:** The intensity of MANF in these brain regions was quantified as described in the *Materials and Methods*. Data were expressed as mean ± SEM, *n* = 6 animals per group. Data were analyzed by Welch’s *t*-test, \**p* < 0.05, \*\**p* < 0.01.

### 3.2. Age-dependent expression of UPR and MANF interacting proteins in the brain

Since MANF play an important role in ER homeostasis and its deficiency exacerbates ER stress in the developing and adult brain [19, 20], we sought to determine whether the observed age-related decline in MANF affects ER stress/UPR in the brain. We examined the expression of UPR proteins during the aging process using IB. As shown in Figs 5 and 6, the expression of PERK declined significantly with age in the cerebellum of both sexes. In the cerebral cortex, the expression of PERK levels was maximal at 4 months and declined thereafter in both sexes. The expression of HYOU1 also decreased with age in the cerebral cortex of both male and female mice. In the cerebellum, HYOU1 expression was highest at 11 months of male mice. In the cerebellum of female mice, HYOU1 expression displayed a trend of downregulation; but the difference did not reach statistical significance (Fig. 6). The levels of phosphorylated-IRE1 (p-IRE1) in the cerebellum significantly declined with age in both sexes, while p-IRE1 levels decreased in male cortex but not females. The expression of GRP94 and GRP78 significantly decreased with age in the cerebral cortex and cerebellum of female mice (Fig. 6). In male mice, their levels remained relatively stable, except for a significant reduction of GRP94 at 4 months in the cerebellum (Fig. 5). In male mice, the expression of ATF6 in the cerebral cortex and cerebellum significantly reduced with age (Fig. 5). In female mice, ATF6 expression in the cerebral cortex declined at 11 months but remained unchanged in the cerebellum (Fig. 6). Interestingly, the expression of ATF4 increased markedly in the cortex of 22-month-old female mice, and a similar elevated trend was observed in male mice. In the cerebellum, ATF4 levels increased at 4 months and then returned to the levels of 1 month in female mice (Figs. 5 and 6). In male mice, the expression of XBP1s was upregulated in both the cerebral cortex and cerebellum with age (Fig. 5). In female mice, the expression of XBP1 in the cortex increased with age, while its levels in the cerebellum decreased (Fig. 6). The levels of phosphorylated-eIF2α (p-eIF2α) in the cortex rose significantly with age in both male and female mice (Figs. 5 and 6). Its levels also increased in the cerebellum of male mice, but not that of female mice. Generally, the levels of total eIF2α in the cortex and cerebellum decreased with age in both sexes (Figs 5 and 6).

**Figure 5.**
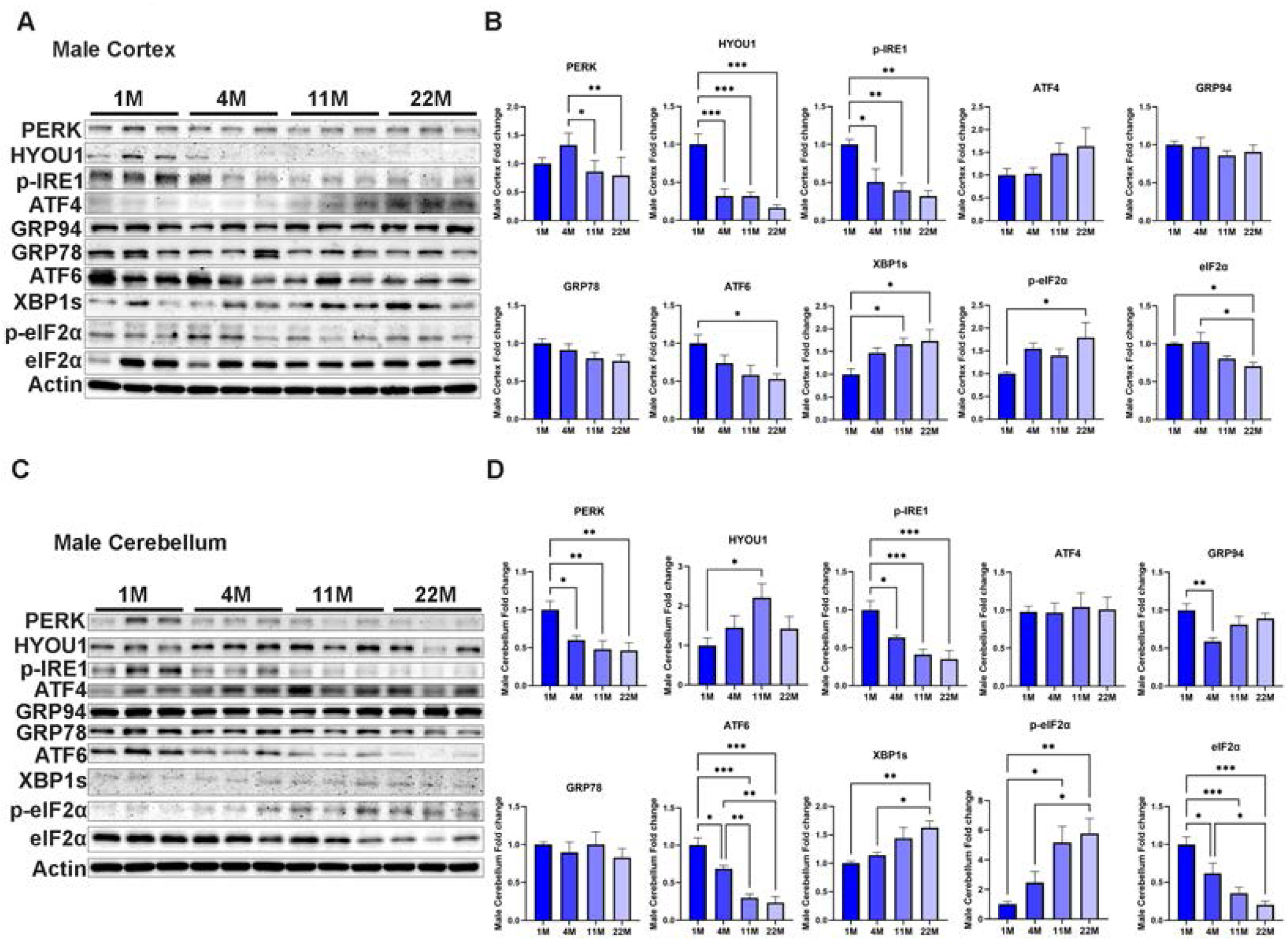
Age-dependent expression of UPR proteins in the brain of male mice. **A** and **C:** Proteins were extracted from the cerebral cortex and cerebellum of 1-, 4-, 11-, and 22- month-old male mice. The expression of UPR proteins was examined by IB analysis. **B** and **D:** The expression of UPR proteins was quantified and normalized to the levels of actin. Data were expressed as mean ± SEM, *n* = 6 per group. Data were analyzed by One-way ANOVA followed by Tukey’s post hoc test, \**p* < 0.05, \*\**p* < 0.01, \*\*\**p* < 0.001.

**Figure 6.**
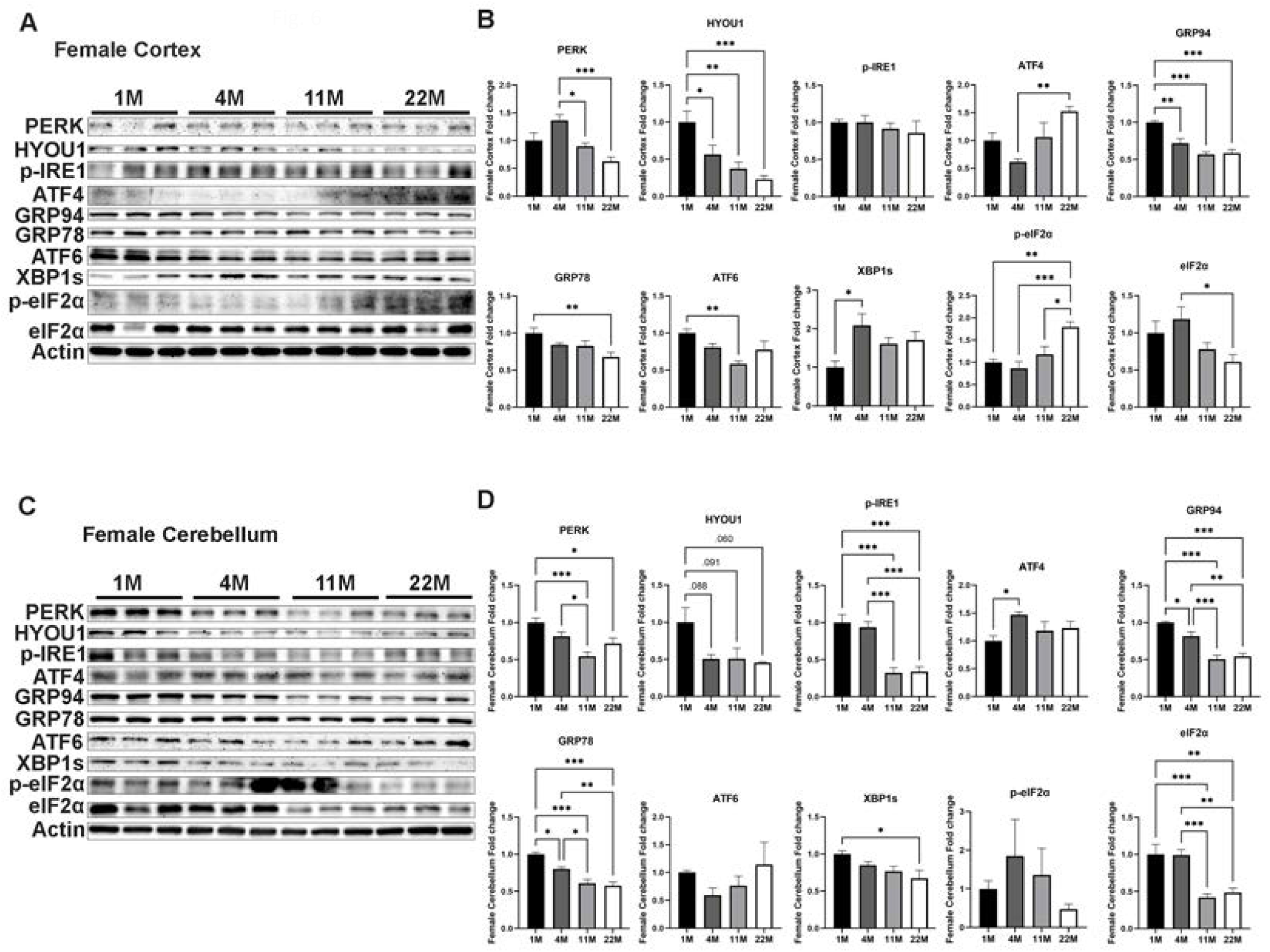
Age-dependent expression of UPR proteins in the brain of female mice. **A** and **C:** Proteins were extracted from the cerebral cortex and cerebellum of 1-, 4-, 11-, and 22- month-old female mice. The expression of UPR proteins was examined by IB analysis. **B** and **D:** The expression of UPR proteins was quantified and normalized to the levels of actin. Data were expressed as mean ± SEM, *n* = 5-6 per group. Data were analyzed by One-way ANOVA followed by Tukey’s post hoc test, \**p* < 0.05, \*\**p* < 0.01, \*\*\**p* < 0.001.

In addition to UPR proteins, the action of MANF may be coordinately mediated by other ER stress-related interacting proteins. Neuroplastin and protein disulfide isomerases (PDIs), including PDIA1 and PDIA6, have been reported to be direct MANF binders and contribute to ER proteostasis and synaptic function [21, 22]. We sought to determine whether the expression of these MANF interacting proteins is also age dependent. In male mice, the expression of neuroplastin, PDIA1 and PDIA6 in the cortex remained unchanged, while the levels of PDIA1 and PDIA6 in the cerebellum declined with age (Figs 7A-D). In female mice, the expression of neuroplastin, PDIA1 and PDIA6 in the cortex declined with age (Figs 7E and F), while the levels of PDIA1 and PDIA6 in the cerebellum decreased with age (Figs 7G and H).

**Figure 7.**
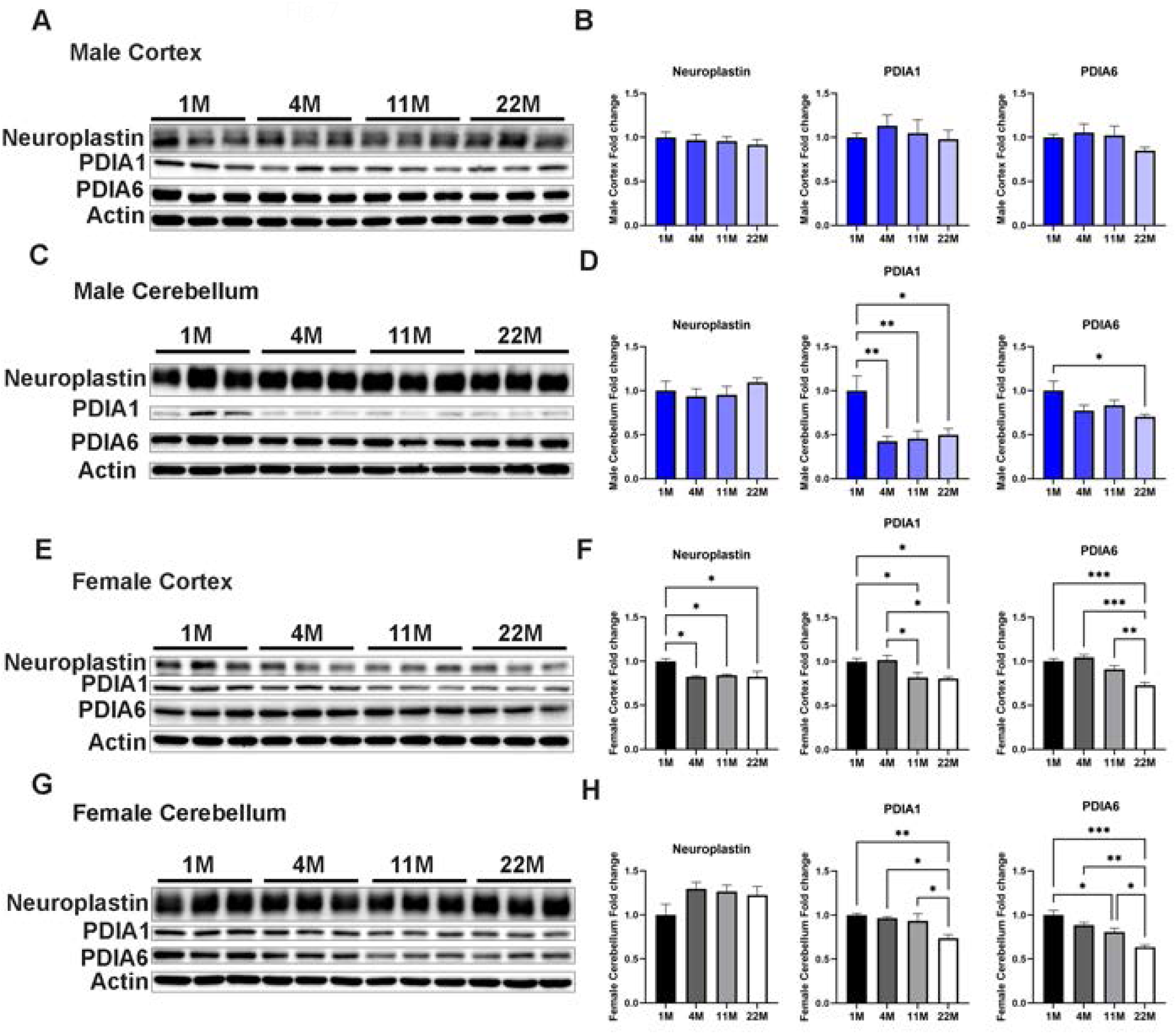
Age-dependent expression of MANF interacting proteins in the brain. **A**, **C**, **E**, **G:** Proteins were extracted from the cerebral cortex and cerebellum of 1-, 4-, 11-, and 22-month-old male and female mice. The expression of MANF interacting proteins (Neuroplastin, PDIA1, and PDIA6) in the cerebral cortex (A) and cerebellum (C) of male mice and that of female mice (E and G) was examined by IB analysis. **B**, **D**, **F**, **H**: The expression of MANF interacting proteins was quantified and normalized to the levels of actin. The results were expressed as mean ± SEM, *n* = 5-6 per group. The data were analyzed by One-way ANOVA followed by Tukey’s post hoc test, \**p* < 0.05, \*\**p* < 0.01, \*\*\**p* < 0.001.

### 3.3. Age-dependent expression of pro-apoptotic and inflammatory proteins in the brain

We sought to determine whether there is an age dependent increase in neuronal death in the brain and examined the levels of cleaved caspase-3 (c-cas-3) and cleaved caspase-12 (c-cas-12) in the cortex and cerebellum of male and female mice (Fig 8). In male mice, there was an age dependent increase of c-cas-3 and c-cas-12 in the cerebellum but not in the cortex (Figs. 8A-D). In female mice, the levels of c-cas-3 but not c-cas-12 in the cortex displayed an age dependent increase (Figs. 8E and F), while both c-cas-3 and c-cas-12 expression in the cerebellum increased with age (Figs. 8G and H).

**Figure 8.**
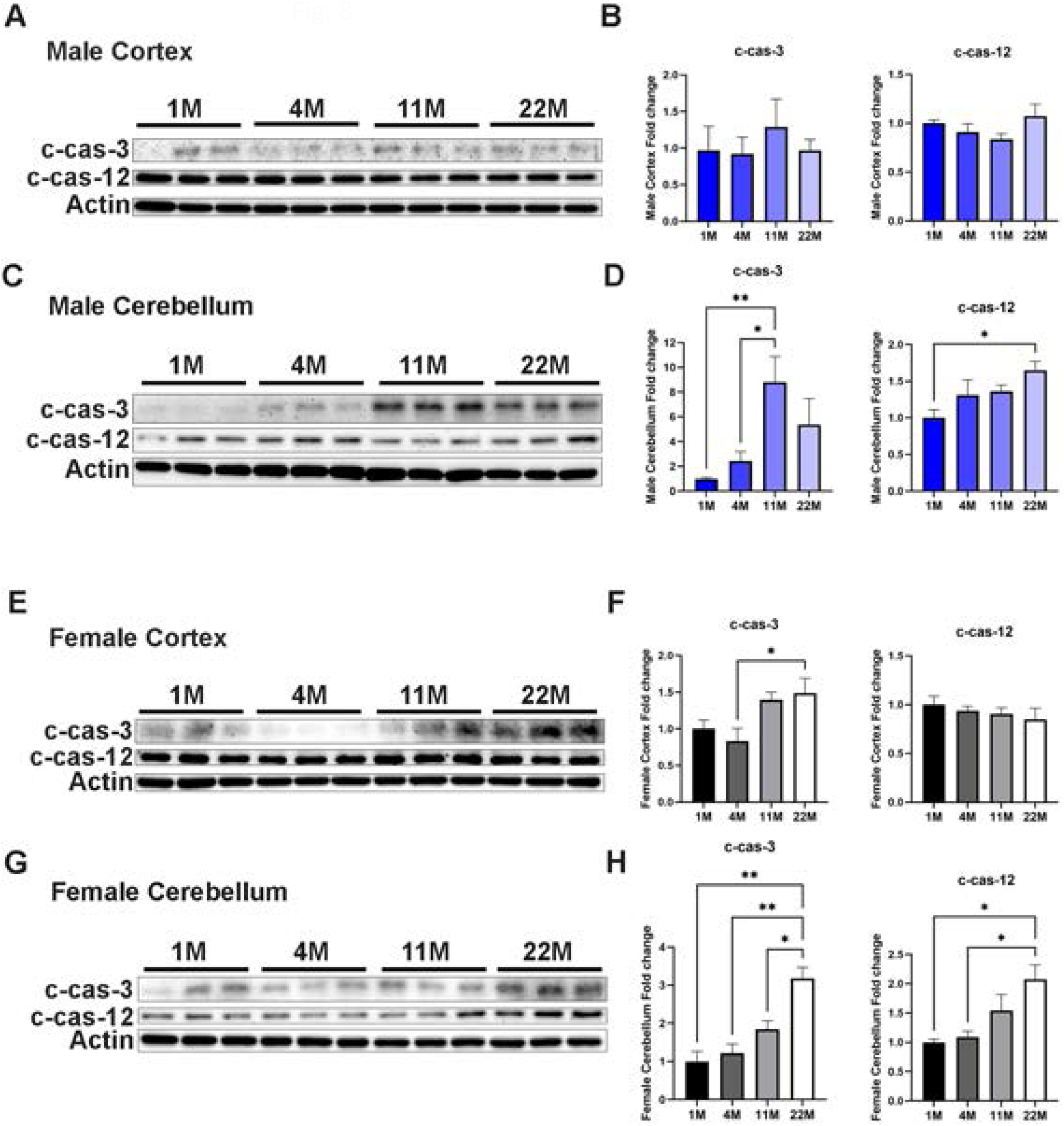
Age-dependent expression of cell death markers in the brain. **A**, **C**, **E**, **G:** Proteins were extracted from the cerebral cortex and cerebellum of 1-, 4-, 11-, and 22-months old male and female mice. The expression of cell death markers (cleaved caspase-3 and cleaved-caspase-12) in the cerebral cortex (A) and cerebellum (C) of male mice and that of female mice (E and G) was examined by IB analysis. **B**, **D**, **F**, **H**: The expression of these cell death markers was quantified and normalized to the levels of actin. The results were expressed as mean ± SEM, *n* = 5-6 per group. The data were analyzed by One-way ANOVA followed by Tukey’s post hoc test, \**p* < 0.05, \*\**p* < 0.01.

To investigate whether there is an age dependent increase of neuroinflammation, we examined the expression of tumor necrosis factor α (TNFα), monocyte chemoattractant protein 1 (MCP1) and ionized calcium-binding adaptor molecule 1 (Iba1) in the cortex and cerebellum of male and female mice across different ages. In the male mice, the expression of TNFα and Iba1 in the cerebellum significantly increased with age (Figs. 9C and D), while only Iba1 levels peaked at 4 month and declined thereafter in the cortex (Figs 9A and B). In female mice, the expression of TNFα in the cortex increased with age (Figs 9E and F), while MCP1 levels peaked at 4 month and declined thereafter in the cerebellum (Figs. 9G and H).

**Figure 9.**
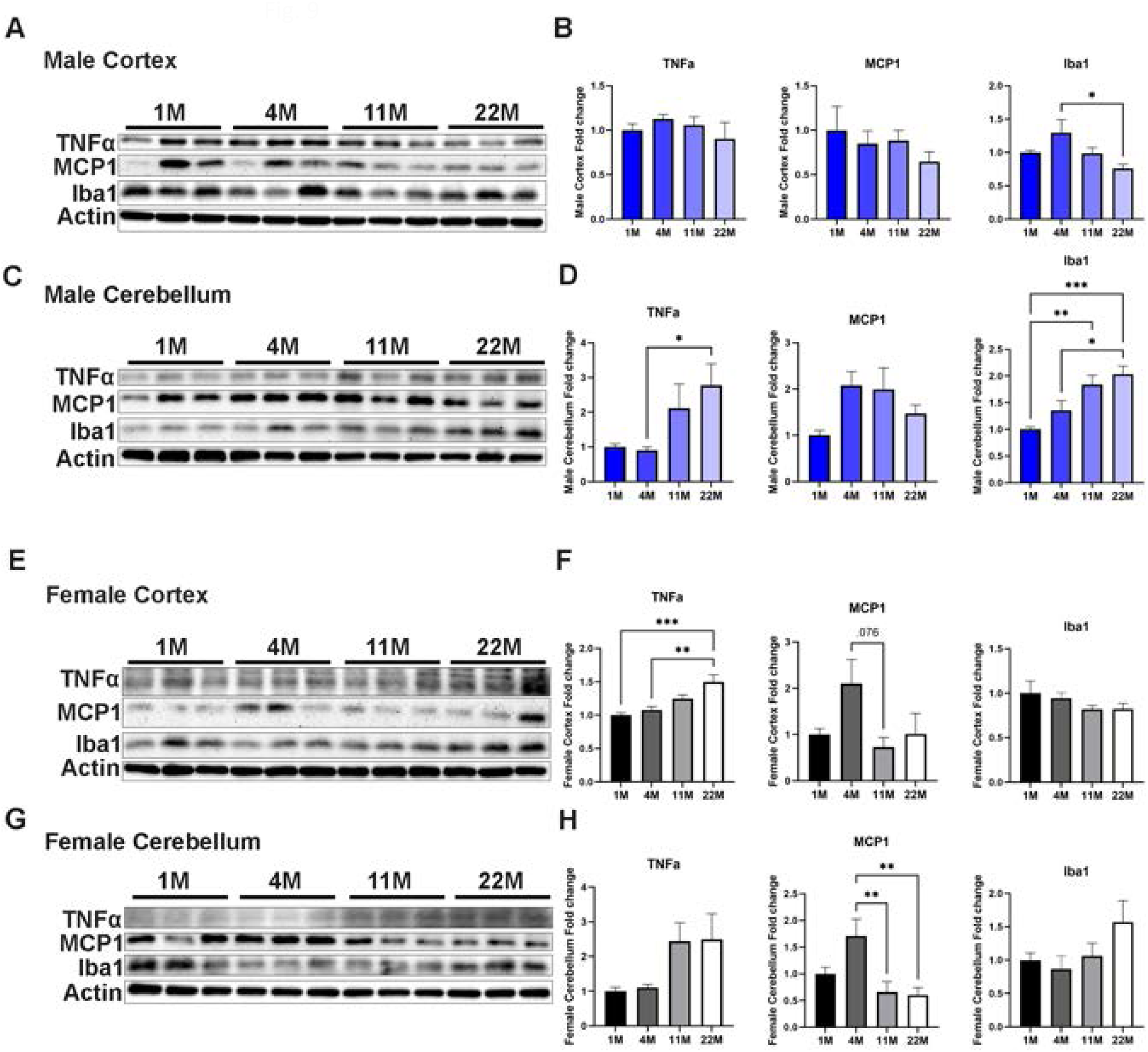
Age-dependent expression neuroinflammatory markers in the brain. **A**, **C**, **E**, **G**: Protein extracts were prepared from the cerebral cortex and cerebellum of 1-, 4-, 11-, and 22-month-old male and female mice. The expression levels of inflammatory markers—TNFα, MCP1, Iba1—were examined by IB analysis. Panels show marker expression in the cerebral cortex (A) and cerebellum (C) of male mice, and in the cerebral cortex (E) and cerebellum (G) of female mice. **B**, **D**, **F**, **H**: Quantification of TNFα, MCP1, Iba1 expression shown in the corresponding panels. Protein levels were normalized to actin and expressed as mean ± SEM, *n* = 5–6 per group. Statistical analysis was performed using one-way ANOVA followed by Tukey’s post hoc test. \**p* < 0.05, \*\**p* < 0.01, \*\*\**p* < 0.001.

### 3.4. Impact of MANF deficiency on neurobehaviors

#### 3.4.1. Generation of cerebellar Purkinje cell (PC)-specific MANF knockout (KO) mice

Since the expression of MANF declines with age, we sought to determine whether MANF deficiency can impact neurobehaviors. Cerebellar PCs had strong expression of MANF which drastically declined with age in both sexes (Figs. 3 and 4). We therefore generated a PC-specific MANF KO mice to evaluate the potential role of MANF in neurobehavioral regulation. As shown in Fig. 10, MANF KO in PCs was confirmed by immunofluorescence (IF) analysis showing the absence of MANF in calbindin-positive PCs of both male and female KO mice at 20 months.

**Figure 10.**
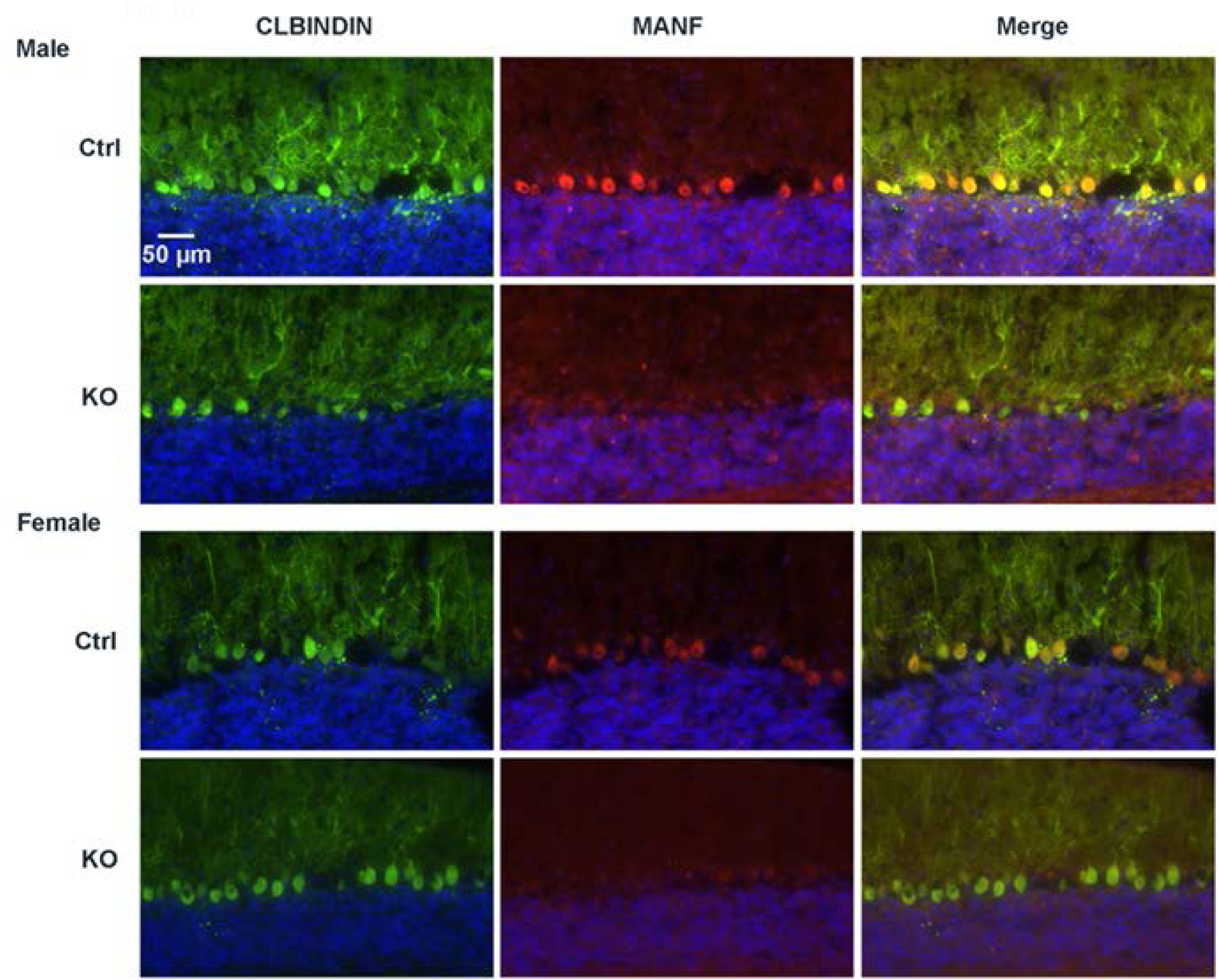
Confirmation of MANF deletion in cerebellar Purkinje cells (PC)-specific MANF knockout (KO) mice. The generation of PC-specific MANF knockout mice was described in the *Materials and Methods*. The loss of MANF expression in PCs was confirmed by immunofluorescence (IF) analysis in 20- month-old control and PC-specific MANF knockout mice. Brain sections were co-labeled with Calbindin (green), a specific marker for PCs, and MANF (red). In control mice, strong MANF immunoreactivity was observed in Calbindin-positive PCs, while in PC-specific MANF knockout mice, MANF expression was absent in PCs, confirming efficient cell-type–specific deletion.

#### 3.4.2. Effects of PC MANF deficiency on anxiety-like behaviors

Open Field test and Elevated Plus Maze test were used to assess anxiety-like behavior in PC MANF KO and control mice of 10-month-old. In Open Field Test, there were no significant genotype differences in total distance traveled or time spent in the center zone for male or female mice (Supplementary Fig. 1). Similarly, in Elevated Plus Maze Test, MANF KO and Ctrl mice did not differ in total distance traveled, percentage of open arm entries, or the percentage of time spent in open arms (Supplementary Fig. 1). These findings indicate that PC MANF deficiency did not affect baseline anxiety-like behaviors.

#### 3.4.3. Effects of PC MANF deficiency on motor coordination

Balance Beam and Rotarod test were used to evaluate motor coordination in PC MANF KO and control mice of 10-month-old. Body weight was measured before the tests. There was no difference of body weight between MANF KO and control male mice; but female KO mice were slightly heavier than control mice (Fig 11A). In Balance Beam test, male MANF KO mice crossed the beams faster than control mice. The difference was statistically significant at 6 mm beam. In contrast, female MANF KO mice required more time to cross the 12 mm and 6 mm beams compared to control mice (Fig. 11B). To determine whether the impaired performance of female KO mice was due to the body weight increase, we conducted correlation analyses between the body weight and the crossing time on the 12 mm and 6 mm beams. The correlation of body weight and crossing time was not significant (Fig. 11C), indicating that the differences observed in motor performance in female MANF KO mice were not due to body weight differences, but rather to MANF deficiency. In Rotarod test, all groups learned to perform better over 5 days training. There were no genotype-dependent differences in male mice. However, the latency to fall was significantly shorter in female MANF KO mice than control mice, indicating impaired motor incoordination in female MANF PC deficiency mice (Fig. 11D). Taken Together, these results suggested a sex-specific effect of MANF deficiency on motor coordination and female mice were more susceptible to MANF deficiency.

**Figure 11.**
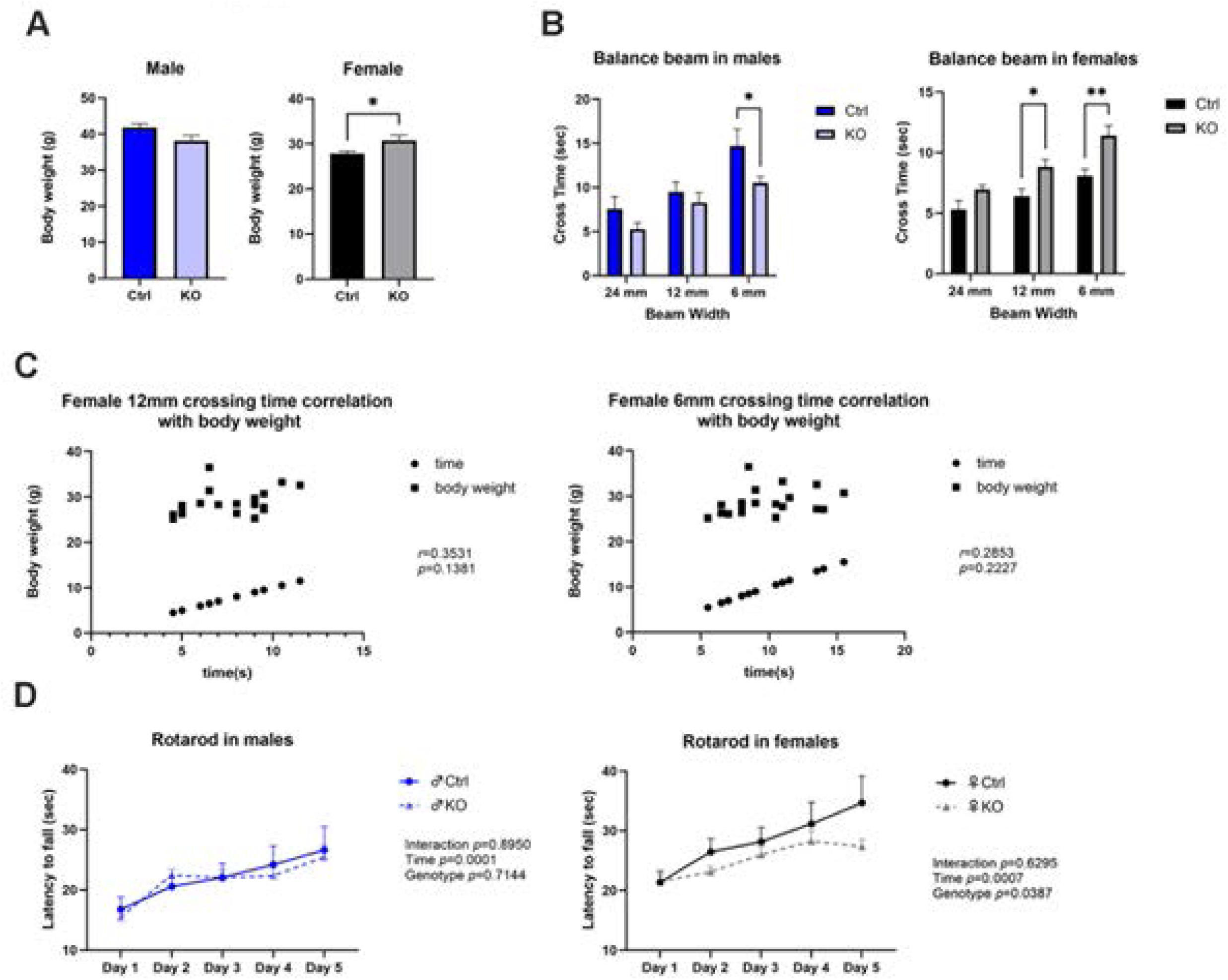
Comparison of motor functions between control and PC-specific MANFKO mice. **A.** Body weight of 10-month-old control and PC-specific MANF KO male and female mice was measured to behavioral testing. **B**. The motor coordination was evaluated using Balance Beam Test. The time taken to cross 24 mm, 12 mm, and 6 mm wide beams was recorded for both male and female mice. **C**. Correlation analysis between body weight and beam-crossing time was conducted for the 12 mm and 6 mm beams in female mice. **D**. Motor performance was further assessed using the Rotarod Test. The latency to fall from an accelerating rotarod was measured over five consecutive days in male and female mice. Data were expressed as mean ± SEM, *n* = 9-11 per group. Panel A and B: Statistical analysis was performed using Welch’s *t*-test. Panel D: Two-way ANOVA was used and followed by Tukey’s post hoc test. \**p* < 0.05, \*\**p* < 0.01.

#### 3.4.4. Effects of PC MANF deficiency on social behaviors

Three-chamber Sociability Task was used to determine social behaviors in PC MANF KO and control mice. Male KO mice appeared more sociable. During the sociability phase, they traveled significantly more distance than control mice (Fig. 12A). They also spent more time in the stranger mouse chamber than in the object chamber (Fig 12B) and engaged in more direct interaction with the social stimulus (Fig 12C). These findings suggested enhanced social motivation and interest in male MANF KO mice. In contrast, female KO mice displayed reduced sociability. They traveled less distances during the sociability test compared to control females (Fig 12D) and spent less time in the stranger mouse chamber (Fig. 12E). Female KO mice also spent significantly less time interacting with the social target compared to Ctrl mice (Fig. 12F). These results revealed a sex-dependent effect of PC MANF deficiency on social behavior, male KO mice display increased sociability, whereas female KO mice exhibit social withdrawal.

**Figure 12.**
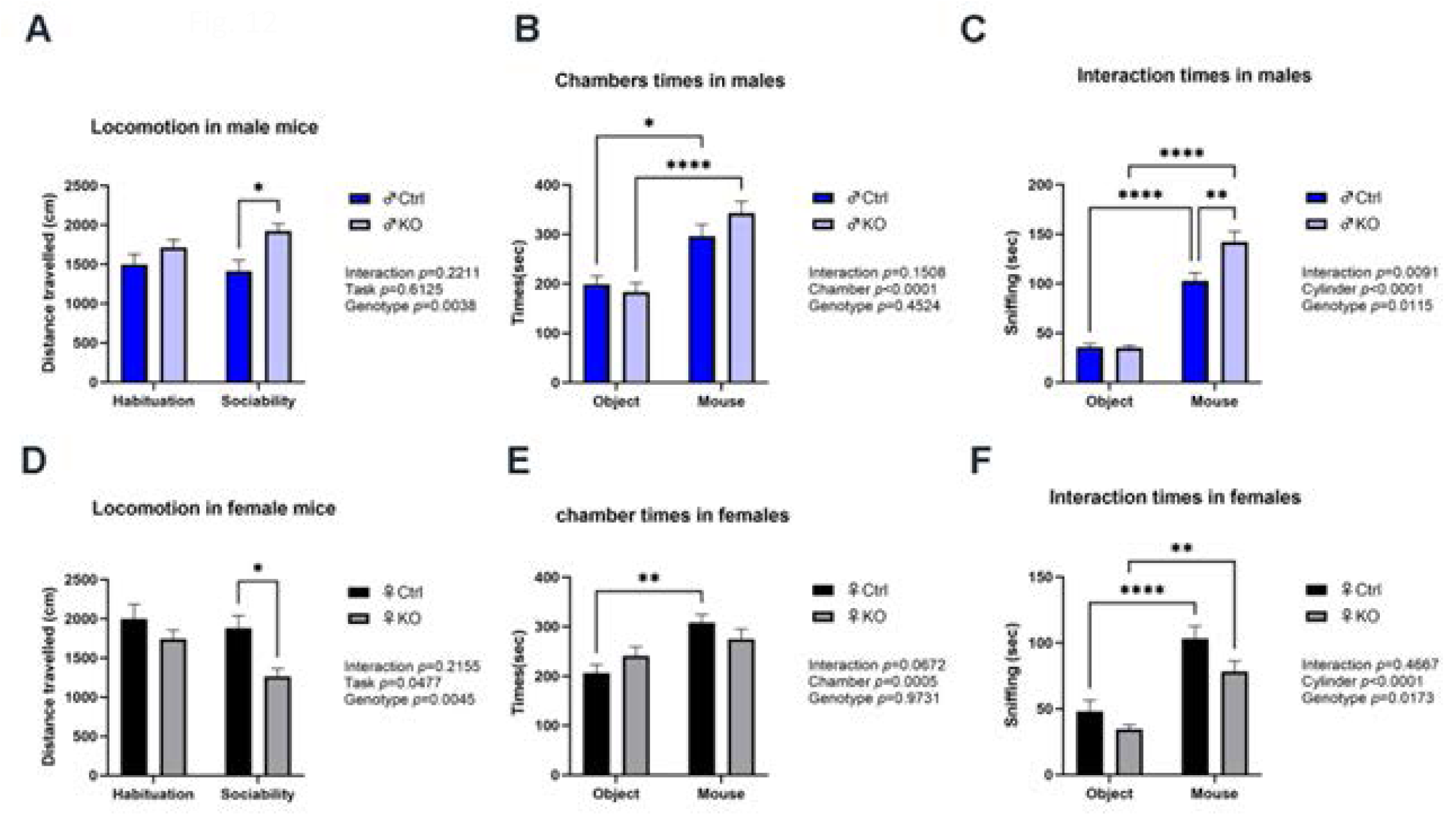
Comparison of social behaviors between control and PC-specific MANF KO mice. Social behaviors were assessed in 10-month-old control and PC MANF KO mice using 3-chamber sociability test. **A** and **D**: Total distance (cm) traveled during the habituation and sociability test periods was measured in male (A) and female (D) mice. **B** and **E**: Cumulative time spent in the chamber with a social mouse versus an object was measured in male (B) and female (E) mice. **C** and **F**: Cumulative time spent interacting with a social mouse versus an object in a cylinder was measured in male (C) and female (F) mice. Data were expressed as mean ± SEM, *n* = 9-11 per group. Data were analyzed by Two-way ANOVA followed by Tukey’s post hoc test. \**p* < 0.05, \*\**p* < 0.01, \*\*\*\**p* < 0.0001.

#### 3.4.5. Effects of PC MANF deficiency on spatial learning and memory

Barnes Maze test was used to evaluate spatial learning and memory (Fig. 13). During acquisition, there was a day effect for both genders, all groups demonstrated learning ability across the 4 days training period (Figs. 13A and D). There was also a genotype effect of male (Fig. 13A) and day vs genotype interactions effect of female (Fig. 13D) in the primary distance traveled to find the escape hole, which demonstrated that both male and female mice MANF KO mice exhibited impaired spatial learning.

**Figure 13.**
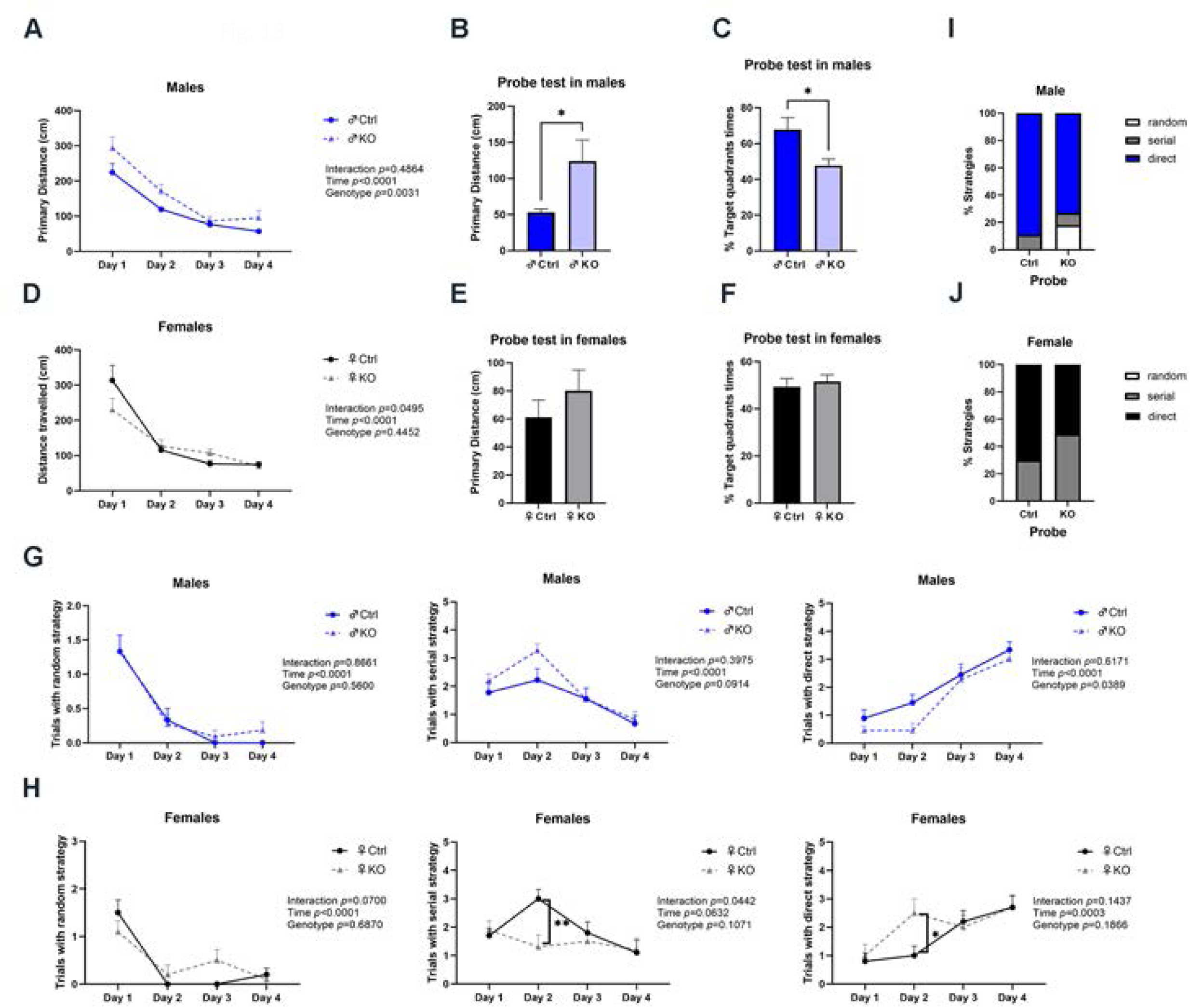
Comparison of learning and memory between control and PC-specific MANF KO mice. Learning and memory were assessed in 10-month-old control and PC MANF KO mice using Barnes maze test. **A** and **D**: Primary distance traveled to locate the escape hole across four days of acquisition was measured in male (A) and female (D) mice. **B** and **E:** Primary distance traveled to locate the escape hole during the probe test was measured in male (B) and female (E) mice. **C** and **F:** Percentage of time spent in target quadrant during the probe test was measured in male (C) and female (F) mice. **G** and **H**: Average number of trials per mouse per day using random, serial, or spatial search strategies across the four days of acquisition was measured in male (G) and female (H) mice. **I** and **J**: Percentage of each search strategy used during the probe test was measured in male (I) and female (J) mice. Data were presented as mean ± SEM, *n* = 9-11 per group. Panel A, D, G and H: Two-way ANOVA followed by Tukey’s post hoc test. Panel B, C, E and F: Welch’s *t*-test. Panel I and J: Fisher’s exact test. \**p* < 0.05, \*\**p* < 0.01.

In the probe trial, which assessed memory retention, male MANF KO mice continued to show memory impairments, traveling longer distances to the target hole (Fig. 13B) and spending significantly less time in the target quadrant (Fig. 13C) compared to control mice. However, female MANF KO mice displayed similar performances as control mice in both primary distances traveled (Fig. 13E) and time spent in the target quadrant (Fig. 13F), indicating that, despite less efficacious learning, their memory was maintained.

Post hoc analysis of searching strategy indicated significant day effects for all strategies in both males and females during the acquisition phase. Male MANF KO mice used less direct (spatial) strategies than control mice (Fig. 13G), while female MANF KO mice showed a significant decreased serial strategy, with a notable day vs genotype interaction effect (Fig. 13H). During the probe test, both male and female KO mice used less direct strategies than control mice (Fig. 13I and J), although these differences did not reach statistical significance. Overall, these findings indicate that PC MANF deficiency may result in impaired spatial learning in both male and female mice, with more robust effects on memory retrieval in males. The reduced preference for efficient strategies in MANF KO mice added to the evidence for MANF involvement in cognition.

## 4. Discussion

In this study, we demonstrated a progressive, age-dependent decline in MANF levels in both the brain and plasma, with the lowest levels observed in old mice (22 months). This reduction in MANF expression was evident across multiple brain regions, including the cerebral cortex, olfactory bulb, thalamus, hypothalamus, hippocampus, and cerebellum. Notably, aged female mice generally exhibited lower MANF levels compared to their male counterparts. Aging also affected the expression of certain UPR proteins as well as proteins that interact with MANF. Using a mouse model with cerebellar Purkinje cell (PC)-specific MANF knockout (KO), we showed that MANF deficiency led to a sex-dependent decline in motor coordination and impacted additional neurobehavioral functions. Collectively, these findings suggest that the age-related decline in MANF expression may contribute to neurological disorders associated with aging and increase vulnerability to neurodegenerative processes.

We examined MANF levels in mice of various ages that range equivalently from early adolescence to the aged in human. A 1-month-old mouse is roughly equivalent to a 10- to 14- year-old human in terms of developmental stages. For 4-month-old mice, the rough human equivalent is about 20–30 years old. 11 months in a mouse is equivalent to 35–40 human years. For 22-month-old mice, the human age equivalent is 65–70 years old. The levels of MANF in the brain and plasma were highest at 1 month and progressively declined to the lowest at 22 months (Figs. 1 and 2). These findings are consistent with a study analyzing MANF levels in human serum [23]. In that study, serum MANF levels were compared among individuals of 21-40 years, 41-60 years and >60 years old. A progressive decline of serum MANF levels with age was observed and lowest levels being observed in > 60 years group [23]. Compared to males, the levels of MANF in the cerebral cortex of females declined more drastically with age (Figs. 1B and D). It appears that the pattern of mRNA expression was different from that of protein levels and there was an increase of mRNA levels at 4 months (Figs. 1E and F). Consistent with the IB data, IHC analysis demonstrated that most of the brain regions examined displayed an age dependent decline of MANF expression in both sexes except the midbrain (Figs 3 and 4). For male mice, CA2 and CA3 of the hippocampus showed no age dependent decrease; while for female mice, it was CA1 and CA3 that showed no age dependent difference. Age dependent decline of MANF expression has been reported in other organs. Sousa-Victor et al compared MANF levels in young (4 months) and old (20 months) mice, and demonstrated reduced levels of MANF in the skin, white adipose tissue, liver and muscle in old mice [23].

An interesting finding from the current study is the presence of sex differences in age dependent expression of MANF and neurobehavioral alterations in MANF deficient mice. Although an age dependent decline of MANF was observed in both sexes, MANF levels in the plasma of female mice were generally lower than male mice (Fig. 2E). Also in the aged cerebellum, MANF expression was significantly less than male mice (Fig. 2D). Furthermore, MANF deficiency in cerebellar PCs affected motor coordination of female mice but not that of male mice (Fig. 11). A substantial body of evidence has accumulated regarding the presence of sex differences in the structure and progression of brain aging [24]. It is currently unclear whether the disparity of MANF expression contributes to aging associated brain impairment. The cerebellum is vulnerable to the aging process with respect to changes of structure, function, and connectivity patterns[25, 26]. Changes in motor coordination and cognitive function associated with cerebellar neuronal dysfunction are among the most important indicators of aging [27]. So far, there is no study investigating sex differences during the process of cerebellum aging. Our findings of reduced MANF levels in aged female cerebellum and increased susceptibility to motor impairment in female PC MANF KO mice may provide an insight into the role of ER stress and UPR in sex differences related to cerebellar dysfunction.

Our study also revealed age dependent alterations in the expression of other UPR proteins. In male mice, some UPR proteins, such as PERK, ATF6, p-IRE1α and eIF2α displayed an age dependent declined in both the cerebral cortex and cerebellum (Fig. 5), suggesting that the brain is losing its capacity to mitigate ER stress during the aging process. However, the levels of p-eIF2α and XBP1s were increased with age, indicative of increased ER stress during the aging. In female mice, more UPR proteins including PERK, GRP94, GRP78, and eIF2α exhibited an age dependent reduction in both the cerebral cortex and cerebellum, and an age dependent decrease of p-IRE1α was only observed in the cerebellum (Fig. 6). The levels of p-eIF2α and XBP1s in the cerebral cortex but not cerebellum increased with age in female mice.

In both sexes, the expression of HYOU1 declined with age except in the cerebellum of male mice (Figs. 5 and 6). HYOU1, also known as 150 kDa Oxygen-Regulated Protein (ORP150) or Glucose-regulated protein 170 (GRP170, is a stress-response protein encoded by the *HYOU1* gene in humans. It is part of the heat shock protein 70 (HSP70) family, which helps protect cells from stress. Although it does not belong to a classic UPR pathway such as PERK, IRE1α, ATF6 and GRP78, HYOU1 acts as a molecular chaperone and primarily resides in the ER [28]. It helps fold newly synthesized proteins and prevents misfolded proteins from aggregating, especially during cellular stress, such as hypoxia, glucose deprivation, and ER stress [28]. HYOU1 has a protective role for neurons and astrocytes in response to cellular stress [29, 30]. Interestingly, HYOU1 is one of MANF interacting proteins which may facilitate MANF’s action [21]. The expression of several other MANF interacting proteins is also affected by the aging process. The most noticeable changes are PDIA1 and PDIA6 which diminished with age in the cerebellum of both male and female mice and their levels also declined in the cerebral cortex of female mice (Fig. 7). PDIA1 and PDIA6 are members of the protein disulfide isomerase (PDI) family that catalyzes the formation, breakage, and rearrangement of disulfide bonds in proteins [31]. They primarily reside in the ER and play crucial roles in protein folding and maintaining ER homeostasis during cell stress, particularly ER stress [31]. PDIA6 also participates in feedback regulation of the UPR by binding and inactivating IRE1α and modulates calcium signaling and apoptosis under prolonged ER stress [32]. PDIA1 and PDIA6 are protective against neurodegenerative diseases [33, 34]. More importantly, PDIA1 and PDIA6 are also MANF interacting proteins and involved in the cellular function of MANF [20, 21]. The diminished expression of PDIA1 and PDIA6 along with MANF with age may further impair proteostasis and promote neurodegeneration. Taken together, the age dependent downregulation of MANF, UPR and MANF interacting proteins implies that the aged brain may be more prone to ER stress, due to the loss of these important proteostasis regulators.

There was an age dependent increase of apoptotic marker, cleaved caspase-3, indicative of injuries and neurodegeneration occurred in the aged brain, particularly in the cerebellum (Fig. 8). The increase of c-cas-3 was accompanied by c-cas-12, suggesting the involvement of ER stress-induced apoptosis, since activation of caspase-12 mediates ER stress-initiated apoptosis.

Cerebellar PCs display a strong expression of MANF which drastically declines with age in both sexes. We used a mouse model of cerebellar PC specific MANF KO mice to evaluate the neurobehavioral outcomes of MANF deficiency. We chose to compare the neurobehavioral profiles of control and PC MANF KO mice at 10 months of age which is equivalent to middle-aged humans, due to the reduced availability of MANF in aged mice. The deficiency of MANF in PCs impaired motor coordination in female mice, which was indicated by poorer performance of MANF KO mice in Balance Beam and Rotarod tests (Fig. 11). PCs are the sole efferent neurons of the cerebellum, sending output from the cerebellar cortex to the cerebral cortex and brain stem, and play a pivotal role in motor and cognitive functions [27, 35, 36]. These results suggest that the reduction of MANF in PCs as a result of aging may impact cerebellar function, impairing motor coordination. It has recently demonstrated that the cerebellum also play an important role in social behaviors [37]. Our findings revealed a sex dependent effect of PC MANF deficiency on social behaviors; male KO mice displayed increased sociability, whereas female KO mice exhibited social withdrawal, suggesting that MANF scarcity in PCs may also impact social behaviors. As discussed above, it is well established that the cerebellum is important in cognition and involved in spatial learning and memory[38]. Our results indicated that PC MANF deficiency caused impaired spatial learning in both male and female mice, with more robust effects on memory retrieval in males (Fig. 13). MANF KO mice showed a reduced preference for efficient searching strategies, supporting that PC MANF is involved in cognition functions. Thus, MANF deficiency in PCs not only impacts motor coordination but also disrupts social behaviors and spatial learning/memory.

Increasing evidence indicates the involvement of MANF in age-related dysfunction. For example, in an inducible mouse model of spinocerebellar ataxia 17 (SCA17), SCA17 disease phenotypes progressed much faster in older mice with earlier neurological symptoms and more severe PC degeneration, along with the reduction of MANF expression [39]. Overexpression of MANF was able to ameliorate neurological symptoms and PC degeneration [39]. MANF was reported to be involved in liver aging and heterozygous mice exhibited progressive liver damage, fibrosis, and steatosis. MANF supplementation reduced several hallmarks of liver aging, prevented hepatosteatosis induced by diet, and improved age-related metabolic dysfunction [23]. It was reported that MANF was associated with an age-related decline in skeletal muscle regenerative capacity [40]. MANF was induced following muscle injury in young mice but not in aged animals, and its expression appeared essential for regenerative success [40]. MANF delivery could improve regenerative impairments in aged muscle in mice [40]. In both mice and humans, retinal aging was associated with a reduction in MANF protein levels, leading to age-related retinal inflammation and increased damage susceptibility [41]. Supplementation of MANF recombinant protein improved retinal homeostasis and repair capacity in aging mice [41]. Ischemic stroke/reperfusion increased MANF expression in brain endothelial cells of aged mice [42]. MANF supplementation could suppress middle cerebral artery occlusion-induced pro-inflammatory factor production, restore brain blood barrier (BBB) integrity and then alleviate infarct volume in aged mice [42]. In conclusion, MANF may play a pivotal role in the aging process of multiple organs. Our findings that the expression of MANF declined with age in the brain and MANF deficiency altered neurobehaviors suggest that targeting MANF may be a practical approach to maintain neuronal functions in aged individuals. We propose in future study to restore MANF levels in aged brain and determine if this is a viable strategy to improve aging associated neurobehavioral deficits.

## CRediT authorship contribution statement

**Di Hu:** Writing – review & editing, Writing – original draft, Methodology, Investigation, Data curation, Formal analysis, Conceptualization. **Wen Wen**: Writing – review & editing, Investigation, Data curation, Conceptualization. **Hui Li:** Writing – review & editing, Methodology, Conceptualization. **Hong Lin:** Writing – review & editing, Project administration, Methodology. **Zuohui Zhang:** Writing – review & editing, Methodology, Conceptualization. **Jia Luo**: Writing – review & editing, Writing – original draft, Visualization, Supervision, Funding acquisition, Investigation, Conceptualization, Project administration.

## Ethics approval

All experimental animal procedures were approved by the Institutional Animal Care and Use Committee (IACUC) at the University of Iowa (#0032295) and performed following regulations for the Care and Use of Laboratory Animals set forth by the National Institutes of Health (NIH) Guide.

## Appendix A.

Supporting information

## Data availability

Data will be made available on request.

## Abbreviations

AD: Alzheimer’s disease
ALS: Amyotrophic Lateral Sclerosis
ARMET: Arginine-rich mutated in early-stage tumors
ATF4: Activating transcription factor 4
ATF6: Activating transcription factor 6
BBB: Brain blood barrier
BSA: Bovine serum albumin
CA1: Cornu ammonis 1
CA2: Cornu ammonis 2
CA3: Cornu ammonis 3
c-cas-3: cleaved caspase-3
c-cas-12: cleaved caspase-12
CNS: Central nervous system
DG: Dentate gyrus
ERAD: ER-associated protein degradation
eIF2α: Eukaryotic Initiation factor 2 alpha
ER: Endoplasmic reticulum
GCs: Granular cells
GRP78: Glucose-regulated protein 78
GRP94: Glucose-regulated protein 94
GRP170: Glucose-regulated protein 170
HYOU1: Hypoxia Up-Regulated Protein 1
HSP70: Heat shock protein 70
IB: Immunoblotting
Iba1: Ionized calcium-binding adaptor molecule 1
IF: immunofluorescence
IHC: Immunohistochemistry
IRE1α: Inositol-requiring kinase 1 alpha
KO: knockout
MANF: Mesencephalic astrocyte-derived neurotrophic factor
MCP1: Monocyte chemoattractant protein 1
NDDs: Neurodegenerative diseases
ORP150: 150 kDa Oxygen-Regulated Protein
PCs: Purkinje cells
PD: Parkinson’s disease
PDI: protein disulfide isomerase
PDIA1: Protein Disulfide-Isomerase A1
PDIA6: Protein Disulfide-Isomerase
A6 PERK: Protein kinase-like ER kinase
SCA17: Spinocerebellar ataxia 17
TNFα: Tumor necrosis factor alpha
UPR: Unfolded protein response
XBP1: X-box binding protein 1

---

**Supplementary Figure 1. Comparison of on anxiety-like behaviors between control and PC-specific MANF KO mice**

Anxiety-like behaviors were assessed in 10-month-old control and PC-specific MANF KO mice using the Open Field Test and Elevated Plus Maze. Open Field Test: **A** and **C**: Total distance traveled during a 10-minute session was measured in male (A) and female (C) mice. **B** and **D:** Percentage of time spent in the center zone during the same session was recorded for male (B) and female (D) mice. Elevated Plus Maze Test: **E** and **H:** Total distance traveled during a 5-minute session was measured in male (E) and female (H) mice. **F** and **I:** Percentage of open arm entries was recorded for male (F) and female (I) mice. **G** and **J**: Percentage of time spent in open arms was recorded for male (G) and female (J) mice. Data were presented as mean ± SEM, *n* = 9-11 per group. Panel A-D: Two-way ANOVA followed by Tukey’s post hoc test. Panel E-J: Welch’s *t*-test.

