## Supplementary figures and images for "Age-Related Alterations in the Expression of Mesencephalic Astrocyte-derived Neurotrophic Factor in the Brain and Their Impact on Neurobehavioral Functions"

### Supplementary Figure 1

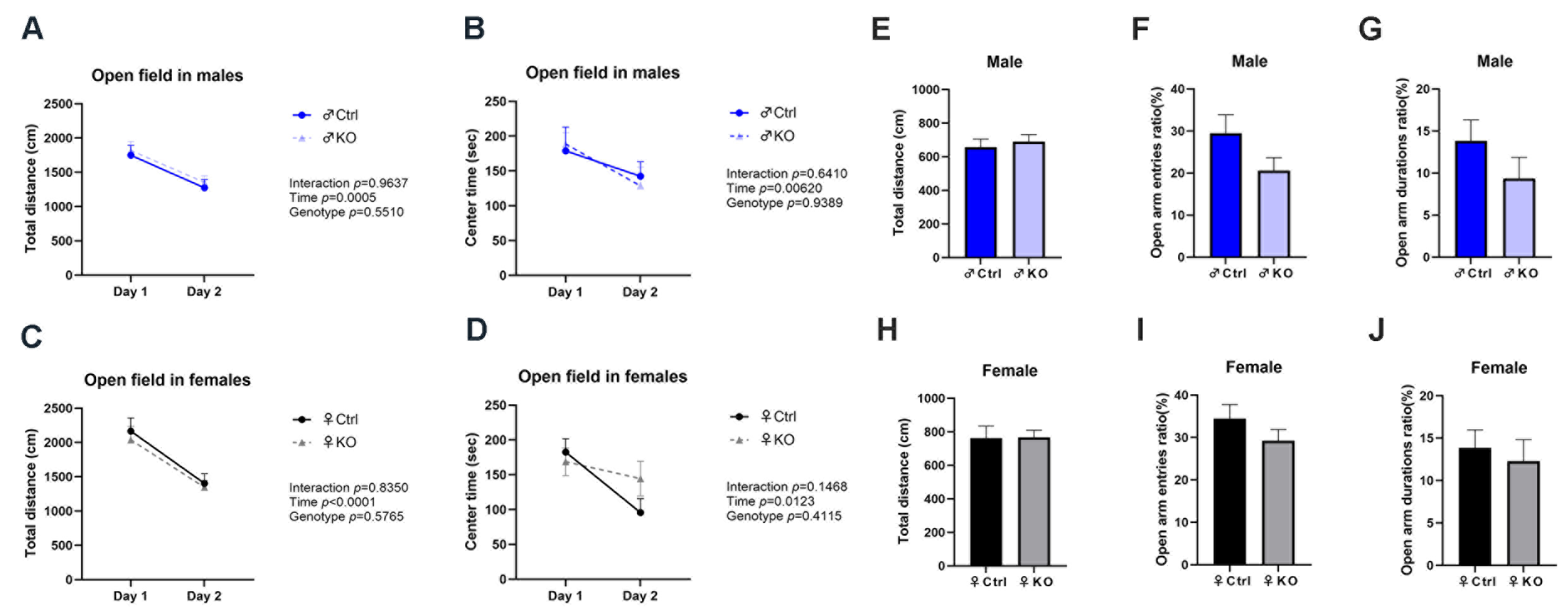
